# Shaping of developmental gradients through selection on multiple loci in *Antirrhinum*

**DOI:** 10.1101/2025.02.12.637738

**Authors:** Desmond Bradley, Louis Boell, Daniel Richardson, Lucy Copsey, Annabel Whibley, Ting Xu, Yu’e Zhang, Yongbiao Xue, David Field, Enrico Coen

## Abstract

Development depends on precise shaping of molecular gradients, but how natural selection acts to establish precision is unknown. Here we analyse genes that control differences in the gradient of yellow flower colour between two varieties of snapdragon (*Antirrhinum*). We show that these differences depend, in part, on *cis* regulatory variation in the pigment biosynthetic gene, *FLAVIA* (*FLA*). *FLA* interacts multiplicatively with three other loci, one of which is a *trans*-acting regulator of *FLA*, to further shape the yellow gradient. All the loci exhibit clines at a hybrid zone, with widths that correlate with phenotypic effect, showing how selection can hone gradient shape with remarkable precision by acting on *cis* and *trans* variation at multiple loci.

## Main Text

Pattern formation is central to development, from establishment of regional identities in early embryos*(1)*, to the generation of colouration patterns in adults*(2)*. A key component of pattern formation is the shaping of gradients, achieved with great reproducibility and precision*(3-5)*. However, it is unclear how natural selection acts to tune gradient properties. Hybrid zones, where species or varieties undergo prolonged gene exchange*(6, 7)*, provide an opportunity to address this problem. Loci under selection can be recognised through steep clines in allele frequency. Some of the identified loci under selection modify colour patterns, such as the aposematic markings on butterfly wings *(8, 9)* or floral guides *(10)*.

Studying how such loci interact can reveal how colour gradients are tuned by selection. Here we apply this approach to flower colour variation between two varieties of snapdragon, *Antirrhinum majus*.

The *Antirrhinum* genus comprises about 25 species primarily distributed throughout the western Mediterranean*(11)*. The species exhibit diverse flower colour patterns. These patterns serve as guides for flower entry by pollinators, mainly bees. Two varieties of *A. majus* subspecies *majus - A*.*m*.*m*. var. *pseudomajus* and *A*.*m*.*m*. var. *striatum (12)* - live in close proximity and have complementary flower colour patterns. *A*.*m*.*m*. var. *pseudomajus* has a yellow highlight (arrowed in Fig. 1A), against a contrasting magenta background; whereas *A*.*m.m*. var. *striatum* has a magenta highlight (arrowed in Fig. 1B) against a yellow background. Yellow and magenta excite different components of the bee visual system, which is based on green-blue-ultraviolet receptors rather than red-green-blue of vertebrates. Yellow excites green receptors, whereas magenta excites blue (ultraviolet reflectance is low for flowers of both varieties*(13)*).

**Fig. 1.**
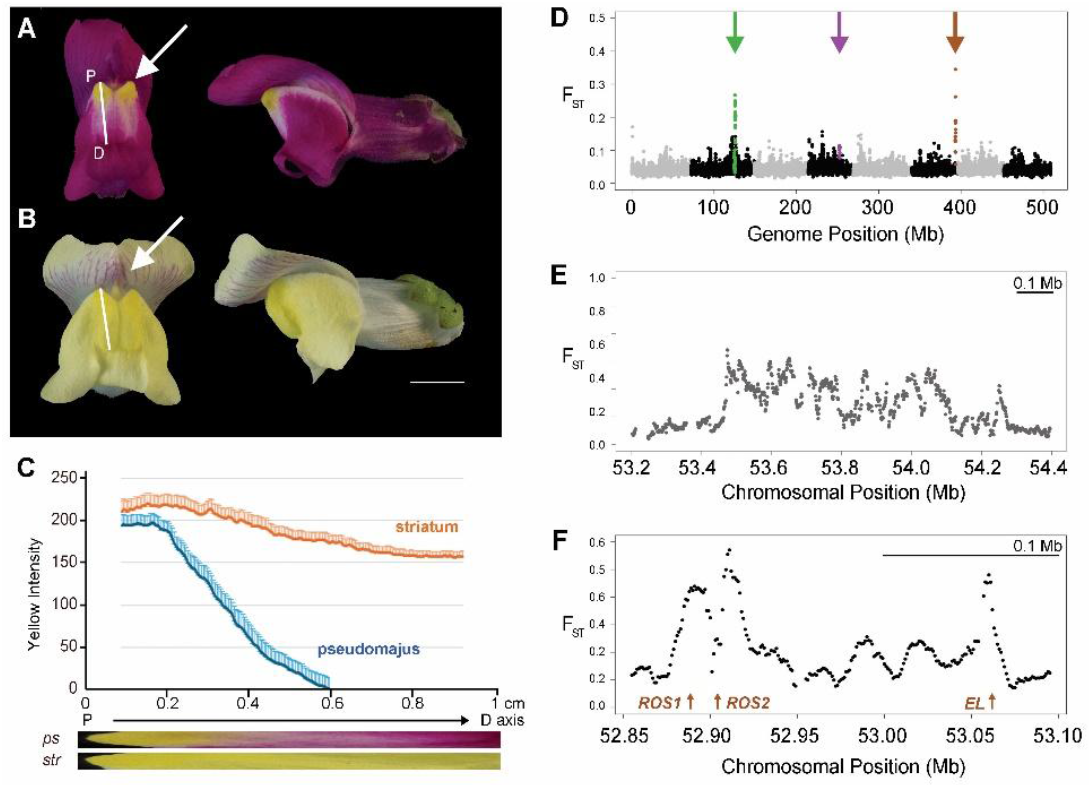
Flower colour phenotypes and *F*_*ST*_ genome scans. (**A**) *A*.*m*.*m*. var. *pseudomajus* flower in face (left) and side (right) views. Arrow indicates yellow highlight (focus) at the bee entry point. White line shows the transect used to measure the yellow gradient, from the proximal (P) to distal (D) edge of the face. (**B**) *A*.*m*.*m*. var. *striatum* flower, with arrow pointing to magenta veins in the upper petals. Scale bar 1 cm. (**C**) Yellow intensity along transects as measured by excess green reflectance over blue for striatum (orange line) and pseudomajus (blue line). For each flower, transect lengths were scaled to 1 cm and reflectance averaged for 10 individuals. Transect photographs are illustrated below the graph. (**D**) Whole genome *F*_*ST*_ scan between a population of *A*.*m*.*m. var. striatum* (YP1) and *A*.*m*.*m*. var. *pseudomajus* (MP4), summarised in windows of 50 kb with 25 kb overlaps. Chromosomes 1 to 8 are shown in alternating black and grey. Purple arrow on chr4 points to the *SULF* locus, brown arrow on chr 6 to the *ROS/EL* loci, and green arrow to a novel island on chr2. (**E**) Close-up of *F*_*ST*_ island on chr2 (10 kb windows, with 9 kb overlaps). (**F**) Close-up of *F*_*ST*_ island on chr6, with *ROS* and *EL* loci denoted by arrows, magnified 4.8 times more than (E) for size comparison.

The yellow colour forms a gradient on the lower (ventral) petal lobe, with the intensity of yellow declining from the point of bee entry towards the edge of the petal. The gradient can be quantified by plotting excess of green over blue reflectance along a transect (white lines in Fig. 1A and B). In *A*.*m*.*m*. var. *pseudomajus*, shows a steep decline in yellow, whereas in *A*.*m*.*m*. var. *striatum*, the yellow gradient is much shallower Fig. 1C). Comparison of four different sets of independent populations gave similar gradient patterns (fig. S1). This difference in yellow gradient steepness is likely related to the distribution of magenta. In *A*.*m*.*m*. var. *pseudomajus*, the steep gradient prevents overlap of yellow with magenta in distal regions of the petal and thus mixing of complementary colours which would reduce visual contrast. In *A*.*m*.*m*. var. *striatum*, magenta is absent in the ventral lobe, allowing extension of yellow into the distal regions of the petal.

Selection on three major loci, *ROSEA (ROS), ELUTA (EL)* and *SULFUREA (SULF)*, contributes to maintaining colour pattern differences between *A*.*m*.*m*. var. *pseudomajus* and *A*.*m*.*m*. var. *striatum* across a hybrid zone *(10, 14, 15)*. In contrast to most of the genome, which shows little differentiation across the hybrid zone, all three loci exhibit steep clines in allele frequency, a signature for barriers to gene flow. *ROS* and *EL* encode *MYB* transcription factors that regulate magenta pigment (anthocyanin) biosynthesis*(16)*; whereas *SULF* encodes small RNAs that restrict the distribution of yellow pigment (aurone) by targeting a biosynthetic gene*(15)*.

*ROS* and *EL* are tightly linked (0.5 centimorgans apart) and exhibit peaks or islands of high relative divergence *(F*_*ST*_) in genome comparisons between populations either side of the hybrid zone (brown arrow, Fig. 1D)*(10)*. *SULF* (purple arrow) does not exhibit a strong *F*_*ST*_ peak, most likely because of polymorphic rearrangements at the locus*(15)*.

### Identification of a new yellow flower locus, *FLAVIA (FLA)*

In addition to the island of high *F*_*ST*_ at *ROS EL*, we observed a second *F*_*ST*_ island (green arrow, Fig. 1D), located on chromosome 2. The chromosome 2 island did not correspond to a known genetic locus responsible for trait differences between *A*.*m*.*m*. var. *striatum* and *A*.*m*.*m*. var. *pseudomajus*, and extended over a greater region than that encompassing the *ROS* and *EL F*_*ST*_ peaks*(10)* (Fig. 1 E,F). To determine whether the greater size of the chromosome 2 island was due to low recombination rates, we genotyped an F2 of *A*.*m*.*m*. var. *striatum* and *A*.*m*.*m*. var. *pseudomajus* for several SNPs in and around the chromosome 2 island (fig. S2A). The recombination rate around the chromosome 2 island was about 0.1 cM/Mb, thirty times lower than 3 cM/Mb measured for *ROS/EL* region*(10)*. Similar low recombination rates were observed when either *A*.*m*.*m*. var. *striatum* or *A*.*m*.*m*. var. *pseudomajus* varieties were crossed to cultivated *A*.*m*.*majus*, indicating that a chromosome inversion between the varieties was not responsible (fig S2B, C).The low recombination rate most likely reflected location of the chromosome 2 island in a pericentromeric region*(17)*.

One explanation for the chromosome 2 island is that it harbours a previously unidentified flower-colour locus under selection. To test this hypothesis, we genotyped 160 plants from a F2 of *A*.*m*.*m*. var. *striatum* crossed to *A*.*m*.*m*. var. *pseudomajus*. We genotyped for a SNP within the chromosome 2 island SNP (SNP2A) as well as SNPs for known colour loci *ROS, EL* and *SULF*. Flowers for each *ROS EL SULF* genotype were ranked visually according to the spread of magenta or yellow flower colour and divided into high (top 50%) and low (bottom 50%) groups (fig S3). All rankings were made blind (before knowledge of genotypes) and repeated three times independently, by different researchers.

No significant differences in SNP2A homozygote frequencies were observed between high and low magenta groups (Fig 2A). However, an excess of *A*.*m*.*m*. var. *striatum* homozygotes and deficit of the *A*.*m*.*m*. var. *pseudomajus* homozygotes was observed in the high versus low yellow group (Fig. 2B). This difference was confirmed by ranking F4 populations segregating for SNP2A in a *ros*^*s*^ *EL*^*s*^ *sulf*^*s*^ background into quartiles (Fig. 2C, fig.S4, allele superscripts refer to origin – *s* for striatum, *p* for pseudomajus). Plants in the lowest yellow quartile were homozygous for the pseudomajus SNP2A allele, whereas those in the highest yellow quartile were homozygous or heterozygous for the striatum SNP2A allele, indicating that the striatum allele was dominant. Similar results were obtained in a *ros*^*s*^ *EL*^*s*^ *SULF*^*p*^ background, though there was greater overlap between quartiles, most likely because reduction in yellow caused by *SULF*^*p*^ made differences harder to discern (Fig. 2D, fig.S5). Thus, the chromosome 2 island harbours a novel yellow flower colour locus, hereafter referred to as *FLAVIA (FLA)*, with the *A*.*m*.*m*. var. *striatum* allele denoted as *FLA*^*s*^ and the *A*.*m*.*m*. var. *pseudomajus* allele *fla*^*P*^.

**Fig. 2.**
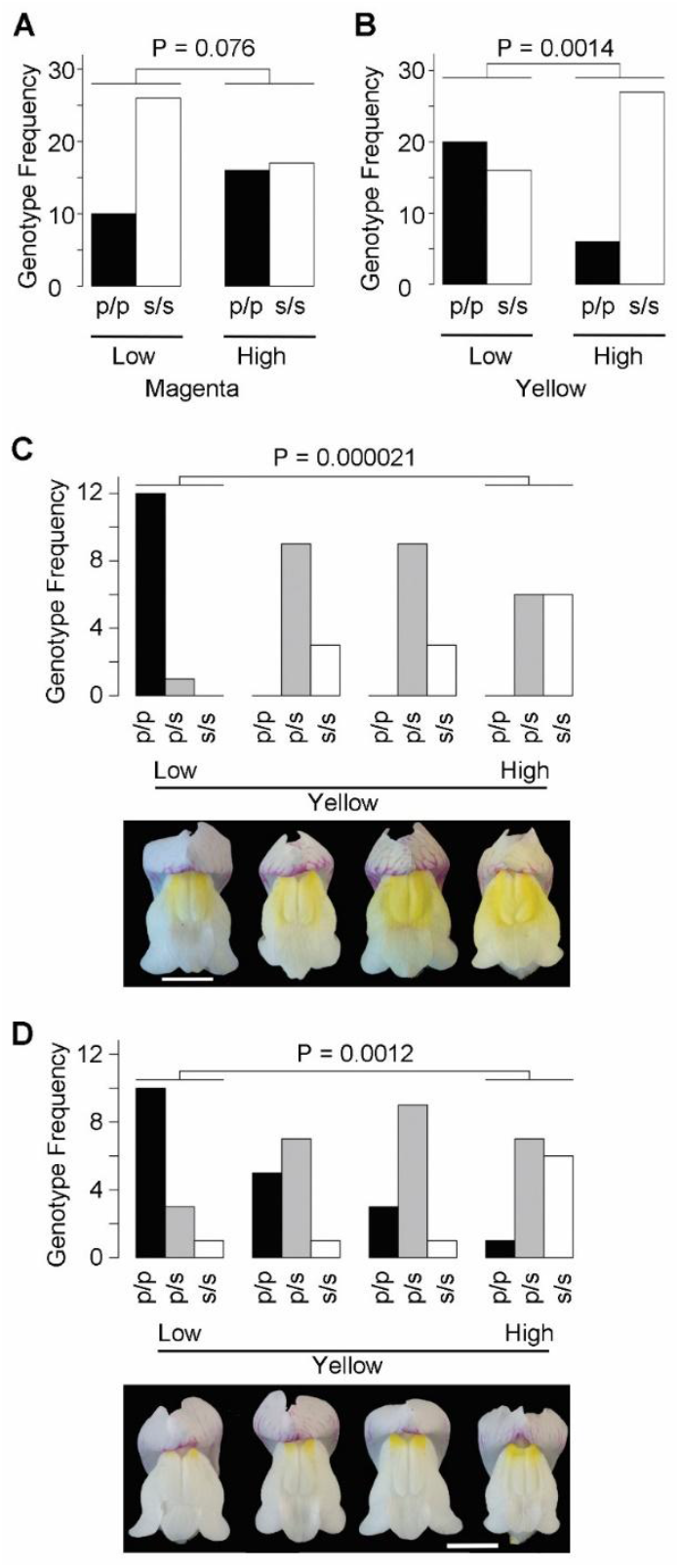
Identification of *FLA*. (**A**) Ranking results from an F2 population (n=69) of *A*.*m*.*m*. var. *pseudomajus (p)* x *A*.*m*.*m*. var. *striatum (s)* ranked for extent of petal lobe magenta and genotyped for SNP2A. Individuals were grouped and ranked separately according to *ROS, EL*, and *SULF* genetic backgrounds, and results were aggregated across all genotypic classes. The low and high magenta categories have no significant difference in frequencies of p/p and s/s homozygotes (P values calculated using a contingency chi-squared test between the low and high quartiles). (**B**) Ranking the same population for extent of yellow shows striatum homozygotes significantly enriched in the high yellow category. (**C**) A *ros*^*s*^ *El*^*s*^*/ros*^*s*^ *El*^*s*^ *sulf*^*s*^*/sulf*^*s*^ population genotyped for SNP2A and ranked for the extent of yellow. The lowest yellow quartile is significantly enriched for p/p homozygotes. Examples of flowers from the middle of each quartile are shown below. Bar is 1 cm. (**D**) A *ros*^*s*^ *El*^*s*^*/ros*^*s*^ *El*^*s*^ *SULF*^*p*^*/SULF*^*p*^ population ranked for their extent of yellow. The lowest yellow quartile is significantly enriched for p/p homozygotes. Bar is 1 cm. Yellow rankings in B and C were done in triplicate by 3 independent observers and all gave similar significant P values (fig.S3).

### *FLA* is the target of *SULF*

To identify the *FLA* gene, we compared RNAseq data from flower buds of *A*.*m*.*m*. var. *striatum* and *A*.*m*.*m*. var. *pseudomajus* for transcripts mapping to the chromosome 2 island (Fig. 3A). The strongest expression difference within the chromosome 2 island (53-fold greater in *A*.*m*.*m*. var. *striatum)* was in transcripts encoding chalcone 4′-O-glucosyltransferase (4’CGT), an enzyme required for aurone (yellow pigment) biosynthesis. 4’CGT is the target of *SULF*, and the low expression in *A*.*m*.*m*. var. *pseudomajus* could therefore be due to repression by *SULF*^*p*^ in *trans*. To test this possibility, we introgressed the chromosome 2 island from each subspecies into a common *sulf*^*s*^*/sulf*^s^ background. The striatum 4’CGT allele was still expressed at a higher level than that of pseudomajus (Fig.3B), indicating that *FLA* likely encodes 4’CGT. Thus, relative to their striatum alleles, *fla*^*p*^ acts in *cis* to reduce *FLA* expression, whereas *SULF*^*p*^ acts in *trans*, via small RNAs, to inhibit *FLA*. RNA *in situ* hybridisations (Fig. 3C, D) showed that *FLA* was expressed in a shallow proximodistal gradient in *A*.*m*.*m*. var. *striatum* but a steep gradient in *A*.*m*.*m*. var. *pseudomajus*, indicating that pseudomajus *cis* and *trans* acting alleles combine to steepen the gradient of *FLA* expression and thus of yellow flower colour.

**Fig. 3.**
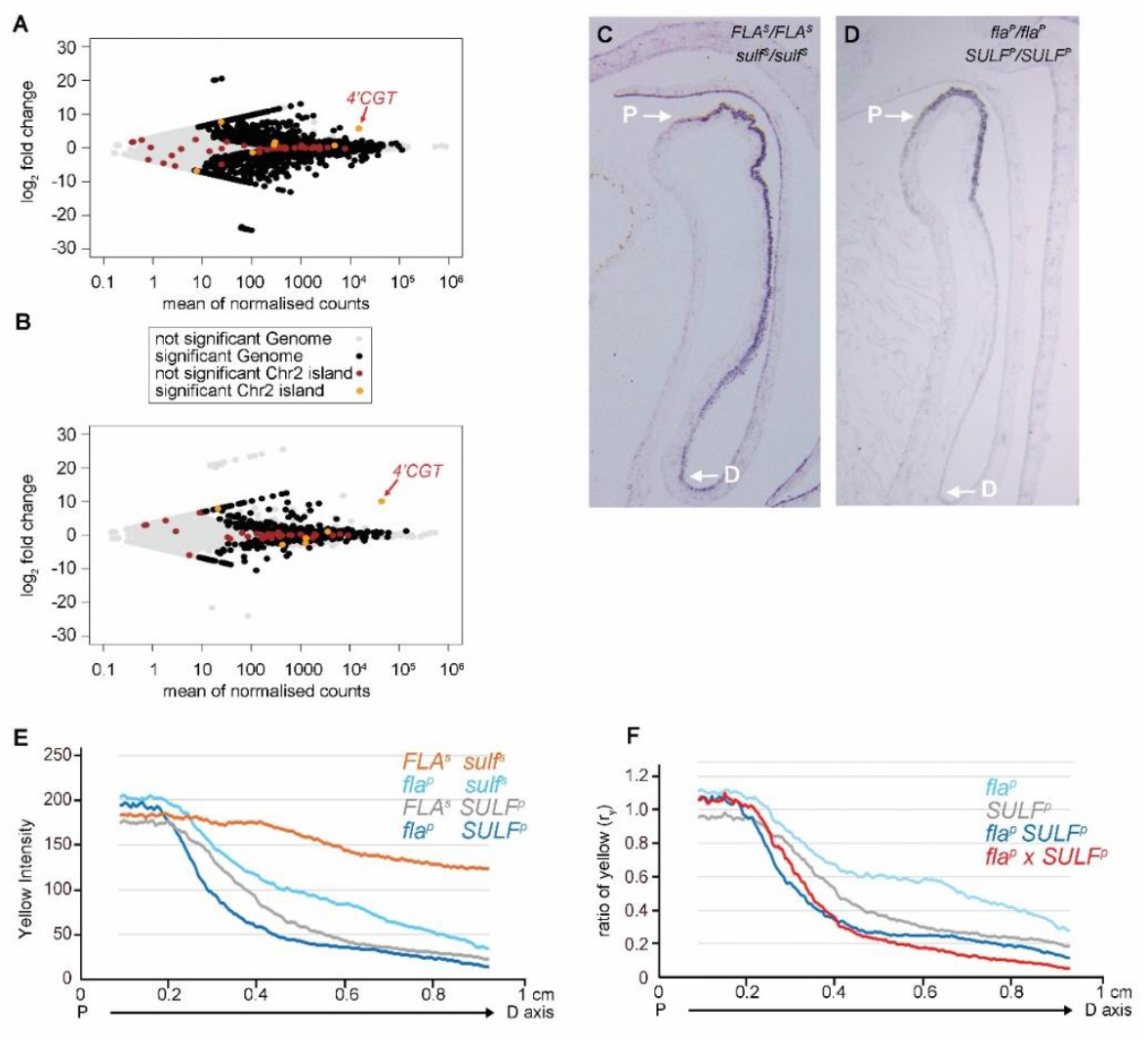
*FLA* likely encodes *4’CGT* and interacts multiplicatively with *SULF*. (**A**) Log fold change in transcripts from the chromosome 2 island between *A*.*m*.*m*. var. *striatum* and *A*.*m*.*m*. var. *pseudomajus. 4’CGT* transcript indicated. Transcripts showing differential expression between the pseudomajus and striatum samples (DESeq2 adjusted p value < 0.01) are highlighted for the genome in black, insignificant in grey. Significant DE transcripts from the Chr2 island are in orange, insignificant island transcripts in brown. (**B**) As in (A) except that the chromosome islands of *A*.*m*.*m*. var. *pseudomajus* and *A*.*m*.*m*. var. *striatum* were introgressed into a common *sulf*^*s*^*/sulf*^*s*^ background. (**C**) RNA *in situ* showing *FLA*^*s*^ expression (dark purple colour) in *A*.*m*.*m*. var. *striatum*. The proximal (P) and distal (D) positions of the petal face used for transect colour profiles are indicated. (**D**) *fla*^*p*^ expression in *A*.*m*.*m*. var. *pseudomajus*. (**E**) Yellow profiles for different combinations of *FLA* and *SULF* alleles (all homozygous), with *FLA*^*s*^ *sulf*^*s*^ (orange) corresponding to the *A*.*m*.*m*. var. *striatum* genotype, and *fla*^*p*^ *SULF*^*p*^ (dark blue) corresponding to the *A*.*m*.*m*. var. *pseudomajus* genotype. Recombinants are in grey or cyan. (**F**) Ratio of yellow intensity, *r*_*y*_, determined by dividing values at each point of the transect in (A) by the value for *sulf*^*s*^ *FLA*^*s*^ at that point. The *fla*^*p*^ x *SULF*^*p*^ predicted *r*_*y*_ (red line) was calculated by multiplying their ry values at each point.

### *FLA* and *SULF* interact multiplicatively to shape the yellow gradient

To understand how *FLA* and *SULF* alleles interact to control the gradient in yellow flower pigment, we compared yellow intensity in transects of the ventral petal lobe, for genotypes in a low magenta *(ros EL)* background (Fig. 3E). Flowers with the *A*.*m*.*m*. var. *striatum* genotype *(FLA*^*s*^ *sulf*^*s*^) had a shallow yellow gradient (orange line). Introducing either *fla*^*p*^ (cyan line, *fla*^*p*^ *sulf*^*s*^) or *SULF*^*p*^ (grey line, *FLA*^*s*^ *SULF*^*p*^), steepened the gradient. Introducing both *fla*^*p*^ and *SULF*^*p*^ (dark blue line, *fla*^*p*^ *SULF*^*p*^), gave an even steeper gradient.

The reduction in pigmentation caused by *A*.*m*.*m*. var. *pseudomajus* alleles could be quantified by calculating the ratio, *r*_*y*_, of yellow intensity of flowers carrying these alleles to that in the *A*.*m*.*m*. var. *striatum* genotype, for each position along the transect. Both *fla*^*p*^ and *SULF*^*p*^ gave a downward proximodistal gradient in *r*_*y*_ showing that pseudomajus alleles acted more strongly to reduce pigmentation at more distal positions (cyan and grey lines, Fig. 3F). Thus, *fla*^*p*^ and *SULF*^*p*^ reduce *FLA* activity in *cis* and *trans* with increasing effect distally. The *SULF*^*p*^ *r*_*y*_ gradient sloped down to lower values than that of *fla*^*p*^, indicating that the *SULF*^*p*^ *trans* effect exceeded the *fla*^*p*^ *cis* effect. The proximodistal gradients in *A*.*m*.*m*. var. *pseudomajus* allele action, as well as the shallow proximodistal gradient of yellow in *A*.*m*.*m*. var. *striatum*, may involve responses to proximodistal prepatterns established earlier in development*(18, 19)*.

A simple gene interaction model posits that for each position in the transect, the combined effect of *fla*^*p*^ and *SULF*^*p*^ on yellow intensity is equal to the product of the effects caused by each locus alone (i.e. the loci interact multiplicatively). Multiplying the *r*_*y*_ values of *SULF*^*p*^ and *fla*^*p*^ together gave a steeper proximodistal gradient (red line, Fig. 3F), similar to the *r*_*y*_ gradient determined experimentally from plants carrying both *SULF*^*p*^ and *fla*^*p*^ (dark blue line). Thus, *SULF*^*p*^ and *fla*^*p*^ alleles interact in a broadly multiplicative manner to steepen the yellow flower colour gradient.

### Selection intensity correlates with effects of *FLA* and *SULF* alleles on the yellow gradient

Both *ROS/EL* and *SULF* exhibit steep clines in allele frequency centred at a similar position at a hybrid zone, with cline widths of about 1km estimated from symmetrical cline fits*(10, 15)*.

Genotyping with SNPs from the *FLA* upstream region revealed a cline at a similar geographical centre though less steep, being about 3 times wider (3km) (Fig. 4A). Cline width is approximately proportional to *σ*/√*s*, where *σ* is the dispersal distance and *s* is the selection coefficient*(9)*. As dispersal distance should be similar for different loci, a threefold greater cline width suggests that the selection coefficient for *FLA* is about 9 times less than for *SULF*. The weaker selection on *FLA* may reflect the smaller effect of *fla*^*p*^ (i.e. higher values of *r*_*y*_) on yellow colour compared to *SULF*^*p*^ (Fig. 3F).

**Fig. 4.**
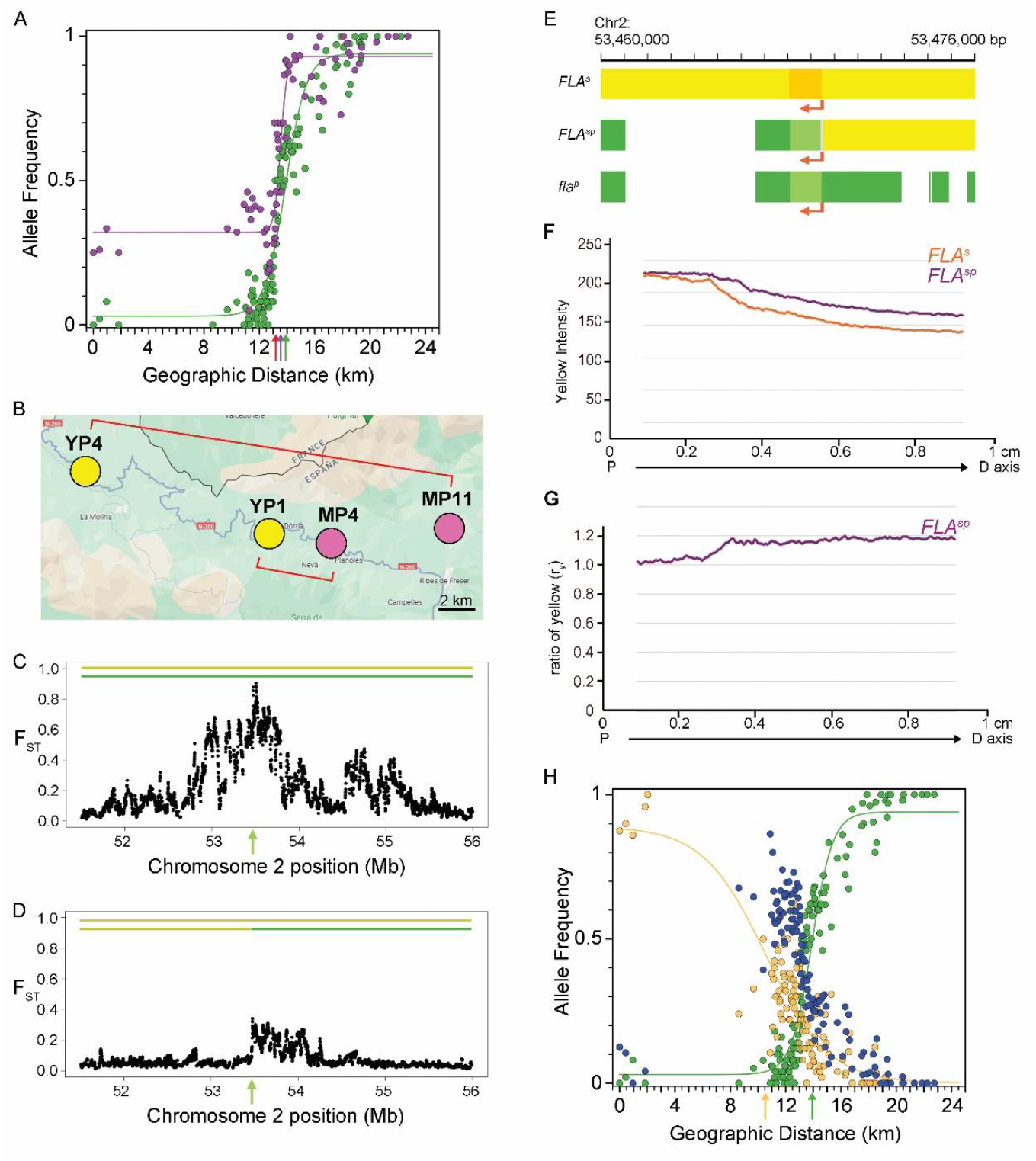
Cline and recombinant analysis. (**A**) SNP allele frequencies indicative of *SULF*^*p*^ (purple points) and *fla*^*p*^ (green points) clines across the hybrid zone (HZ) from *A*.*m*.*m*. var. *striatum* (west) to *A*.*m*.*m*. var. *pseudomajus* (east) flanking populations. Based on sigmoid fits, the cline widths are about 1km for *SULF*^*p*^ and 3km for *fla*^*p*^, with cline centres marked by arrows (including ROS in red). The non-zero frequency of the *SULF* pseudomajus SNP on the *A*.*m*.*m*. var. *striatum* flank is likely caused by one or more *sulf*^*s*^ alleles carrying the pseudomajus SNP (through recombination or rearrangement). **(B)** Location of DNA pools analysed either side of the hybrid zone. (**C**) *FLA F*_*ST*_ comparison with distant populations YP4 and MP11. The major alleles present in the comparison are shown by the coloured lines; *FLA*^*s*^ (yellow) and *fla*^*p*^ (green). The FLA coding region is indicated by a green arrow. (**D**) *FLA F*_*ST*_ comparison with near populations YP1 and MP4. Lines show the major alleles; *FLA*^*s*^ (yellow) and *FLA*^*sp*^ (yellow-green) with the FLA coding region indicated by the green arrow. (**E**) Schematic showing the genomic structures of the 3 different *FLA* alleles found in the HZ. *FLA*^*s*^ (yellow), *FLA*^*sp*^ (5’region in yellow, breakpoint in grey, coding in light green, and 3’ region in darker green), and *fla*^*p*^ (green). Transcription direction shown by arrows. Indels are shown as white gaps relative to the reference *A*.*m*.*majus. FLA*^*sp*^ has the 5’ region of *FLA*^*s*^ joined to the coding and 3’ of *fla*^*P*^. (**F**) Yellow profile of face region for the *FLA*^*s*^ and *FLA*^*sp*^ alleles. (**G**) Ratio of yellow intensity, *r*_*y*_, of the *FLA*^*sp*^ allele vs *FLA*^*S*^ allele. (**H**) *FLA*^*s*^ (yellow), *fla*^*p*^ (green) and *FLA*^*sp*^ (purple) allele frequencies in demes across the HZ, with clines fitted for *FLA*^*s*^ and *fla*^*p*^. Cline centres are shown by arrows and widths are about 3km for *fla*^*p*^ and 9km for *FLA*^*s*^.

The relationship between phenotypic effect and fitness could be further investigated through analysis of a recombinant *FLA* allele found to be prevalent in the hybrid zone. Comparisons of populations further away from the hybrid zone centre (populations YP4 and MP11, Fig. 4B) gave a broader and taller *F*_*ST*_ island around *FLA* (Fig. 4C) than comparisons between nearby populations (YP11 and MP4, Fig. 4D). SNP analysis revealed that this difference was accounted for by a recombinant allele, hereafter named *FLA*^*sp*^, at a high frequency near the centre of the hybrid zone. *FLA*^*sp*^ had the upstream region derived from *A*.*m*.*m*. var. *striatum* and the downstream region derived from *A*.*m*.*m*. var. *pseudomajus* (Fig. 4E). The recombination breakpoint was located between a SNP at -64 and +40 relative to ATG start codon of *FLA* (fig. S6). For distant population comparisons, the predominant allele on the *A*.*m*.*m*. var. *striatum* side was *FLA*^*s*^, giving a broad *F*_*ST*_ island; whereas in the closer populations, the predominant allele on the *A*.*m*.*m*. var. *striatum* side was *FLA*^*sp*^, truncating the island after the recombination breakpoint.

Analysis of petal transects showed that *FLA*^*sp*^ gave a similar gradient in yellow to *FLA*^*s*^ but slightly shallower (Fig. 4F). The steeper yellow gradient conferred by *fla*^*p*^ compared to these alleles (Fig. 3E) therefore largely reflects its *cis-*acting upstream region (absent in *FLA*^*sp*^ and *FLA*^*s*^). The *r*_*y*_ values for *FLA*^*sp*^ were greater than 1 and sloped upwards along the proximodistal axis, showing that *FLA*^*sp*^ boosts *FLA* expression slightly relative to *FLA*^*s*^ distally (Fig.4G). Thus, the downstream region of *FLA*^*s*^ (absent in *FLA*^*sp*^) contains *cis*-acting elements that attenuate expression in distal petal positions.

Based on the above analysis, we used SNPs from various positions in the *FLA* locus to infer frequencies for three alleles: *fla*^*p*^, *FLA*^*s*^ and *FLA*^*sp*^ (Fig. 4H). The *FLA*^*s*^ cline (yellow) was about three times wider than the *fla*^*p*^ cline (9km compared to 3km), indicating that selection on *FLA*^*s*^ was about 9 times lower than on *fla*^*p*^. *FLA*^*sp*^ (purple) showed a peak in frequency just to the left of the *fla*^*p*^ cline centre.

The high frequency of the *FLA*^*sp*^ allele near the centre of the hybrid zone most likely reflects its selective advantage in a hybrid genetic background. The hybrid zone is at least 100 years old and may be considerably older*(10)*. Suppose initial contact led to the formation of complementary frequency clines for *FLA*^*s*^ and *fla*^*p*^, with geographic centres coincident with the clines of magenta-controlling loci *ROS EL*. Many plants on the left side of the cline centre would have yellow flowers with overlapping weak magenta caused by introgression of *ROS*^*p*^ *el*^*p*^ alleles. In this context, a recombinant allele at *FLA* that enhanced yellow, *FLA*^*sp*^, may have had a selective advantage over parental *FLA*^*s*^ by over-riding the dulling effect of magenta. *FLA*^*sp*^ would then have been swept to high frequency in this geographical location. Outside of this location, *FLA*^*sp*^ may have lower fitness because it confers excess yellow (on *A*.*m*.*m*. var. *striatum* side) the or a gradient that is too flat (on *A*.*m*.*m*. var. *pseudomajus* side) to guide pollinators effectively.

This selective sweep would have led to subdivision of the hybrid zone into a sequence of three regions, each with a different predominant *FLA* allele: a left region with high *FLA*^*s*^ frequency, a middle region with high *FLA*^*sp*^, and a right region with high *fla*^*p*^. Selection, and thus cline width, would reflect fitness differences between *FLA* alleles in adjacent regions. The large cline width for *FLA*^*s*^ (9km) would correspond to a small fitness difference between *FLA*^*s*^ (left region) and *FLA*^*sp*^ (middle region), correlating with a small phenotypic effect of *FLA*^*sp*^ relative to *FLA*^*s*^ (Fig. 4F). The narrower cline width for *fla*^*p*^ (3km) would correspond to a greater fitness difference between *FLA*^*sp*^ (middle region) and *fla*^*p*^ (right region), correlating with a greater phenotypic effect of *fla*^*p*^ relative to *FLA*^*sp*^ (Fig. 3E, 4F). This cline width is greater than that of *SULF*^*p*^ (1km) which has a stronger phenotypic effect than *fla*^*p*^ (Fig. 3F). Thus, selection coefficient broadly correlates with the strength of phenotypic effect, which likely reflects the salience of visual signals to pollinators.

### Two additional loci shape the yellow gradient through multiplicative interactions

Recently, three additional loci controlling flower colour differences between *striatum* and *pseudomajus* were identified by genome scans for clines*(20)* and phylogenetic signatures*(21)*: one affecting magenta, *RUBIA (RUB)*, and two affecting yellow, *AURINA (AUN)* and *CREMOSA (CRE)*. These loci have weaker phenotypic effects than the *ROS EL* and *SULF* loci, and do not exhibit strong *F*_*ST*_ peaks. *AUN* likely encodes Aureusidin synthase (AS1), which catalyses the step before 4’CGT in the aurone synthesis pathway*(22)*.

Yellow profiles of *AUN* and *CRE* in genetic backgrounds fixed at the other loci showed that both *A*.*m*.*m*. var. *pseudomajus* alleles reduce yellow (Fig. 5A and B, fig.S7,S8). Values of *r*_*y*_, gave a slight negative slope, indicating that pseudomajus alleles steepened the gradient in yellow but to a lesser extent than *FLA* or *SULF* (Fig.5C), at least in the genetic backgrounds analysed.

**Fig. 5.**
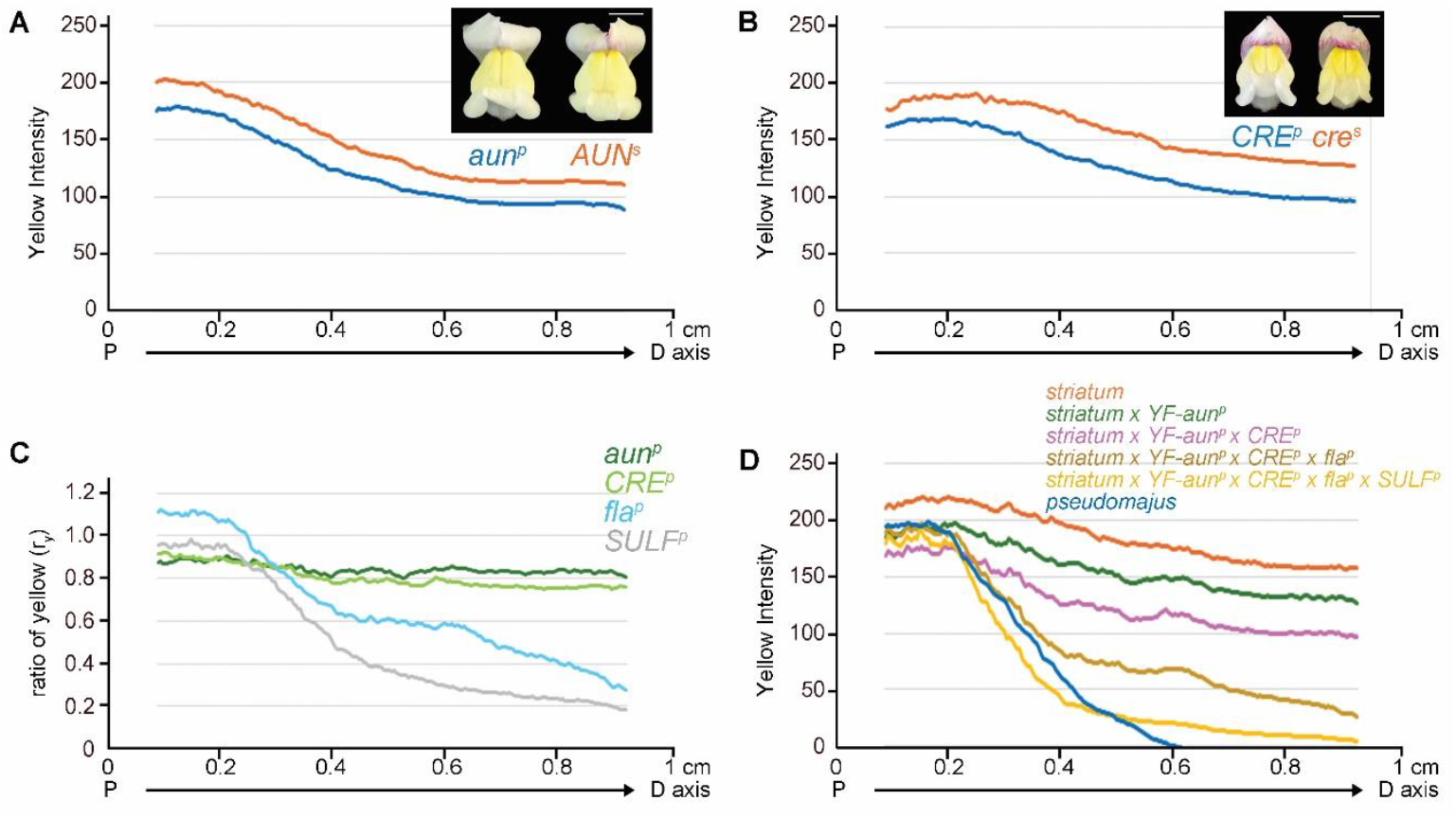
Effects of *AUN* and *CRE* and multiplicative model. (**A**) Yellow profile of face region for the *AUN*^*s*^ and *aun*^*p*^ alleles in a *sulf*^*s*^*/sulf*^*s*^ *ros*^*s*^*/ros*^*s*^ *FLA*^*s*^*/FLA*^*s*^ *CRE*^*p*^*/CRE*^*p*^ background Example flowers shown. **(B)** Yellow profiles for the *cre*^*s*^ and *CRE*^*p*^ alleles in a *sulf*^*s*^*/sulf*^*s*^ *ros*^*s*^*/ros*^*s*^ *FLA*^*s*^*/fla*^*p*^ *AUN*^*s*^*/aun*^*p*^ background. Example flowers shown. (**C**) Ratio of yellow *(r*_*y*_) of the *aun*^*p*^ vs *AUN*^*s*^ allele, and *CRE*^*p*^ vs *cre*^*s*^, with *fla*^*p*^ vs *FLA*^*s*^ and *SULF*^*p*^ vs *sulf*^*s*^ shown for comparison. (**D**) Gradient in yellow obtained by multiplying the *A*.*m*.*m*. var. *striatum* profile (orange) by *r*_*y*_ values of different loci in combination. *A*.*m*.*m*. var. *pseudomajus* profile shown in blue for comparison with the profile obtained by multiplying the effects of all four loci together (yellow).

To determine how the four yellow loci, *AUN, CRE, FLA* and *SULF*, interact, we assumed a multiplicative model (i.e. multiplying their values of *r*_*y*_ together) and calculated the yellow intensity they would confer by progressively introducing more *A*.*m*.*m*. var. *pseudomajus* alleles into *A*.*m*.*m*. var. *striatum* (Fig. 5D). Introduction of *A*.*m*.*m*. var. *pseudomajus* alleles progressively steepened the gradient of yellow reducing yellow in the most distal position shown (0.92 cm) by 97% when all four were combined. The gradient of yellow in the proximal half for all four alleles combined was similar to that observed for the *A*.*m*.*m*. var. *pseudomajus* flowers (blue line). In the distal half, the gradient of yellow in *A*.*m*.*m*. var. *pseudomajus* could not be accurately estimated because of the confounding effect of magenta.

The steep gradient in yellow colour in *A*.*m*.*m*. var. *pseudomajus* is therefore established through multiplicative interactions between four loci. The effect of single alleles at some loci (e.g. *AUN* and *CRE)* is hardly perceptible to the human eye, but collectively their effects multiply to produce a steep gradient. Our findings are consistent with genome-wide surveys in yeast, which have shown that functional gene interactions are most commonly multiplicative*(23)*.

For the observed clines in allele frequency to be maintained, selection must prevent alleles at each locus from introgressing from one subspecies into the other. There are no obvious differences in physical environment or pollinators between *A*.*m*.*m*. var. *striatum* and *A*.*m*.*m*. var. *pseudomajus* populations*(24, 25)*. Selection therefore most likely maintains the clines because of the epistatic interactions between loci controlling magenta *(ROS, EL, RUB)* and yellow *(SULF, FLA, AUN, CRE)* flower colour. In *A*.*m*.*m*. var. *pseudomajus*, the seven loci interact to produce a flower with a yellow highlight against a largely non-overlapping a magenta background. An allele that flattens the yellow gradient, causing it to overlap with magenta would muddy the colours making the flower less attractive to pollinators. In *A*.*m*.*m*. var. *striatum*, the seven loci interact to produce a flower with a magenta highlight on a graded yellow background. An allele that further steepens or flattens the yellow gradient in this context may reduce pollinator attractiveness or guidance. In a genetic background where alleles at loci have been mixed, as in the centre of the hybrid zone, alleles such as the recombinant, *FLA*^*sp*^, can be favoured over parental ones.

## Conclusion

Our results indicate that honing of the yellow developmental gradient has been shaped with great precision through selection acting on multiple loci rather than through modifications at a single locus, consistent with the notion of evolutionary tinkering*(26)*. The phenotypic effect of individual alleles can be very small yet confer fitness differences sufficient for selection to maintain steep clines between hybridising populations. Fitness differences depend on epistatic interactions with other loci, notably those controlling magenta colour. Thus, depending on genetic context, different gradients may be favoured.

Precise shaping of developmental gradients has been described for other systems, such as the Bicoid morphogen gradient in the *Drosophila* egg*(4)*. Unlike the yellow gradient in *Antirrhinum*, the Bicoid gradient is established through diffusion and dynamic control of morphogen degradation, rather than in response to a prepattern. It is not known how selection has tuned the Bicoid gradient but it may have involved alleles that slightly shift, steepen or flatten the gradient by modifying production, diffusion or degradation parameters, which may vary spatially across the embryo*(27)*. Some of these alleles may have been at Bicoid, but others may have been at different loci controlling these parameters. Thus, selection on multiple loci, each with alleles having modest effects, may provide a general mechanism for evolutionary shaping of developmental gradients.

## Acknowledgements

We thank Catherine Taylor for plant care, Nick Barton for sharing SNP data and useful comments, Hugo Tavares for useful discussions and bioinformatics, Matt Couchman for field and data archiving, Tingting Li for help with photography and phenotyping, Jordi Chan for help with ImageJ analyses, and Xana Rebocho for organisation of field experiments.

## Author’s contributions

Conceptualization: LB DF EC

Data Curation: DR AW DF EC

Formal analysis: DR AW DF EC

Funding acquisition: YX NB DF EC

Investigation: DB LB DR LC

Methodology: LB DR AW DF EC

Project administration: YX DF EC

Resources: DR DF

Software: DR AW DF

Supervision: YX DF EC

Validation: DR DF

Visualization: DB DR DF EC

Project administration: DF EC

Writing – original draft: EC

Writing – review & editing: DB DR AW DF EC

## Data and materials availability

Raw DNA and RNA data have been uploaded to SRA (accession number SUB15081578). The *A. majus* reference genome V3.0 is available at the NGDC Genome Warehouse (accession number GWHBJVT00000000).

## List of Supplementary Materials

Materials and Methods

Figs. S1 to S9

Tables S1

References (28-31)

