## Supplementary Data for "Shaping of developmental gradients through selection on multiple loci in *Antirrhinum*"

**SUPPLEMENTARY MATERIALS**

Materials and Methods

Figs. S1 to S9

Table S1

References (28–31)

**Materials and Methods**

**Plant Materials**

The locations of the *Antirrhinum* species accessions used in this study were mapped (fig. S9). An interactive location resource is found at: <http://antspec.org/>

Seed from Accessions and John Innes stocks were grown on soil-compost mixes in pots or trays, outside on benches in the summer, or in the greenhouse with supplemental light in winter as described (28). Lines from self-incompatible species were maintained by inter-sibling crossing.

*Antirrhinum majus majus* (*FLA^m^/FLA^m^; SULF^m^/SULF^m^; ROS^m^ el^m^/ROS^m^ el^m^*) John Innes reference line JI7 was used to make the Snapdragon Genome Reference Sequence (<http://bioinfo.sibs.ac.cn/Am/> ) and used in crosses for studying *FLA^m^* alleles.

*A.m.m.* var. *pseudomajus* (*fla^p^/fla^p^; SULF^p^/SULF^p^; ROS^p^ el^p^/ROS^p^ el^p^*) reference line seeds were collected from allopatric populations near Ventola ~5km east of the Pyrenees Hybrid Zone.

*A.m.m.* var. *striatum* (*FLA^s^/FLA^s^*; *sulf^s^/sulf^s^*; *ros^s^ EL^s^/ros^s^ EL^s^*) reference line seeds were collected from allopatric populations near La Molina ~ 10 km west of the Pyrenees Hybrid Zone. Self-incompatible wild *Antirrhinum* lines were maintained by intercrossing siblings at each generation.

Wild Accession pools of *A.m.m.* var. *pseudomajus* or *A.m.m.* var. *striatum* were harvested from ~30-50 individuals from different locations as described (9, 20, 21).*A.m.m.* var. *pseudomajus* and *A.m.m.* var. *striatum* Accessions used were also maintained from wild collected seed by intercrossing siblings and their original locations mapped (fig.S9).

F2s and introgressions.

*A.m.m.* var. *pseudomajus* x *A.m.m.* var. *striatum* F2:

*A.m.m.* var. *pseudomajus* from Ventola plant J1428 seed capsule 2, was sown to give individual V163-36. *A.m.m.* var. *striatum* from La Molina plant J1324 seed capsule 5, was sown to give individual V206-40. V163-36 was crossed with V206-40 to generate F1 plants (Y132 -1 to 5) which were self-incompatible and therefore sibs were intercrossed to give the F2 family J109.

*A.m.m.* var. *striatum* x *A.m. majus*:

*A.m.m.* var. *striatum* from La Molina plant J1160 seed capsule 3 was sown to give individual V189-16. *A.m. majus* JI7 was crossed with V189-16 to generate F1 plants (Y137) that were self-pollinated to give F2 family J106.

*A.m.m.* var. *pseudomajus* x *A.m. majus*:

*A.m.m.* var. *pseudomajus* came from Ventola plant J428 seed capsule 2, which was sown to give individual V163-36. *A.m. majus* JI7 was crossed with V163-36 to generate F1 plants (Y135) that were self-pollinated to give F2 family J108.

F3s and F4s.

For Fig.2C, an F4 *sulf^s^/sulf^s^* population was generated from intercrossing individuals of the *A.m.m.* var. *pseudomajus* x *A.m.m.* var. *striatum* F2 family J109 above to give F3 families K115 and K116. Individuals from these families were crossed to give F4 family L116 (*ros^s^ EL^s^/ros^s^ EL^s^; sulf^s^/sulf^s^*) segregating for *FLA^s^* and *fla^p^*.

For Fig.2D, an F4 *SULF^p^/SULF^p^* population was generated from intercrossing individuals of the *A.m.m.* var. *pseudomajus* x *A.m.m.* var. *striatum* F2 family J109 above to give F3 family K113. Two sibs from this family were intercrossed to give F4 family V162 (*SULF^p^/SULF^p^*) segregating for *ROS^p^ el^p^* and *ros^s^ EL^s^*^,^ and for *FLA^s^* and *fla^p^*. For ease of scoring, only *ros^s^ EL^s^/ros^s^ EL^s^* individuals were analysed.

To obtain all 4 combinations of *SULF^p^* vs *sulf^s^* and *FLA^s^* vs *fla^p^* for analysis in Fig.3, we used *ros^s^ EL^s^/ros^s^ EL^s^* individuals families L116 and V162 families above. They were further genotyped and selected as fixed for *cre^s^/CRE^p^* and *AUN^s^/aun^p^*, but segregating for *FLA^s^* and *fla^p^*, before selecting 10 individuals for analysis.

The effects of other loci analysed in Fig.5, used the populations L116 for *CRE* and V164 for *AUN*. Genotyping of family L116 identified *sulf^s^/sulf^s^; ros^s^ EL^s^/ros^s^ EL^s^; FLA^s^/FLA^s^*; *cre^s^/cre^s^* individuals segregating for *AUN^s^* and *aun^s^*. Family V164 provided individuals with genotype *sulf^s^/sulf^s^; ros^s^ EL^s^/ros^s^ EL^s^; FLA^s^/fla^p^; AUN^s^/aun^p^* segregating for *cre^s^* and *CRE^p^*. Again, 10 individuals with the different *CRE* genotypes were colour profile analysed.

**Leaf genomic DNA Isolation**

For genotyping, 3-6 small young leaves (~1cm long; total 100-200mg) were collected in Eppendorf tubes and frozen in liquid nitrogen. Large numbers were collected in 96 tube or plate format and frozen at -20^o^C before isolation by the QIAGEN DNeasy 96 Plant Kit or an in-house Genotyping Service method as described in: [Wheat and Barley DNA Extraction in 96-well Plates - MASWheat (yumpu.com)](https://www.yumpu.com/en/document/view/11880064/wheat-and-barley-dna-extraction-in-96-well-plates-maswheat#google_vignette).

For genomic DNA, samples from the greenhouse were stored at -80^o^C until extraction. Leaf samples collected from field locations in France and Spain were either placed in bags in silica, or stored in moist paper towel and kept cool at 4^o^C until they could be Courier Posted by overnight delivery to the lab in the UK and frozen at -80^o^C on arrival. Genomic DNA was isolated by a CTAB method using ~2-10g of leaves harvested either from a single individual or as pooled samples as described (29).

**Photography**

Flowers were placed on black velvet with a scale bar and Small Grey & Colour Separation Chart (Danes Picta BST13) for colour, light level and white balance monitoring. We used an Olympus XZ-1 (10 Megapixels) Camera with side/overhead lighting via table lamps fitted with halogen 42W 630 lumen (2800k) warm white light bulbs. Camera settings were set to the closest White Balance of 3000K, no *fla*sh, Macro On, F stop 8.0, Exposure Time 1/20-1/40 sec, ISO 200, RAW images, aspect 4:3, high definition.

**Phenotyping**

Yellow and magenta patterns were scored for flowers taken fresh from each individual plant. Scores were made by eye using the numerical system as described (13). Photographs of each flower, face and side views were taken for reference and re-checking. Flowers were also cut in half to give lower ventral/lateral petals separated from upper/dorsal. These allowed optimal imaging of yellow colour in the face region as used in ImageJ

**ImageJ analysis of flower colour profiles**

10 different individual flower photographs were selected of the lower half of each flower; foci to the edge of petal lobes. Each original jpg was first adjusted in Photoshop using -

Image-adjustments-levels – to standardise the white balance, using the white reference colour chart square included in all photographs. Images were collected in one Photoshop image at 600 dpi RGB and all backgrounds made black.

Each individual was adjusted to the same size of 1 cm from foci to the most distal point on the petal face. Flowers were then orientated so that this foci to distal point transect was vertical. The image was saved as a jpg and opened in Image J.

In ImageJ a 1 pixel width line was drawn to run through the vertical foci to the edge of the petal lobe. The flower image was split into the 3 colour channels and the blue, green and red profiles collected in excel. We calculated yellow as green pixels minus blue pixels. Fractional effects (FE) were calculated from the yellow profile (YP) at each point along the transect of the petal face: ratio of yellow r_y_ = YP of pseudomajus value/YP of striatum value. This allowed us to calculate the effect of any pseudomajus allele on any striatum YP. For example, the r_y_ for each locus, *FLA*, SULF, AUN and CRE could be used against the YP of an *A.m.m.* var. *striatum* to see how close it makes the profile to an *A.m.m.* var. *pseudomaju*s profile.

**F_ST_ Analysis**

The comparison of population pools for F_ST_ determination was as described (21).

**Measuring Chr2 Recombination Rate**

Families were generated (see ‘Plant Materials’ above) that allowed recombination between *A.m.m.* var. *pseudomajus* and *A.m.m.* var. *striatum* (J109), *A.m.m.* var. *pseudomajus* and *A.m. majus* (J108) or *A.m.m.* var. *striatum* and *A.m. majus* (J106). Each F2 family was genotyped with KASP markers (Supplemental table S1) along the chromosome. The resulting genotypes and recombination events were used to calculate recombination frequences along chr2, based on the number of gametes screened to give Supplemental fig.S2.

**Genotyping Plants**

The KASP Genotyping method (LGC) was as described (14) and the fluorescence signals discriminating homozygous or heterozygous alleles were detected by a BioRad CFX96 Lightcycler and the data processed using the BioRad CFX Manager software v3.1.

The KASP method (LGC Genomics) was used to identify *FLA*, *SULF*, *AUN* and *CRE* alleles using the oligo sets listed in Supplemental table S1 below.

AFLP markers were also used to genotype the EL locus and confirm *CRE* families (table S1). Standard PCR conditions were used, with annealing at 55^o^C and extension steps at 72^o^C of 1min. PCR products were run on 1% (w/v) agarose gels, stained with ethidium bromide (~0.5 ug/mL) and photographed under UV light.

**Flower Colour Ranking**

Flower photographs included the Danes Picta BST13 colour chart, allowing all flower images to be white-balanced equivalently in Photoshop. By eye, each image was assessed for the yellow spread/intensity and positioned within a ranking order. Ranking order was continually updated as more flowers were analysed. The final ranking was divided into four quartiles. Within each quartile set, the *FLA* genotypes were determined for each individual, and the number of *A.m.m.* var. *pseudomajus (p)* and *A.m.m.* var. *striatum (s)* alleles were recorded.

**RNAseq**

Total RNA was isolated from petal tissues using the Qiagen RNeasy Plant Mini Kit, including DNaseI treatment. For a comparison of *A.m.m.* var. *pseudomajus* (accession Ac1266) and *A.m.m.* var. *striatum* (accession Ac1125) we harvested petal lobes from dissected flowers just before opening. Three independent samples were used for reproducibility, and each sample was a pool of 3 individuals, each contributing 1-2 flowers. Samples were sent to the Earlham Institute, Norwich, UK for inhouse library and Illumina sequencing runs to give 2x75bp Paired-End sequencing of strand-specific libraries.

For a comparison of *sulf^s^/sulf^s^ FLA^s^/FLA^s^* with *sulf^s^/sulf^s^ fla^p^/fla^p^* (in the segregating F2 population J109), only upper/dorsal petal lobes were used. As above, triplicate samples were analysed as described above. These samples were sent to Novogene, Cambridge, UK and processed as above.

All RNAseq data was processed, mapped and analysed as described (21)

**RNA *in situ***

*FLA* probes and method were as described (30). In addition, pre-*in situ* photographs were taken of the wax sections as the flower pigments had been preserved and thus highlighted the patterns of yellow, magenta and non-pigmented regions. Pigmentation was subsequently lost on dewaxing and processing of tissue for *in situ*.

**Geographic Cline Analysis**

**Plant sampling and SNP genotyping**

At the Planoles hybrid zone, Los Pironeos, Spain, individual plants were sampled for leaf material and flowers (colour phenotyping) and geolocated (with Trimble GEOX) as part of another study building a pedigree at the site over generations. These samples came from surveys over a period of 6 to 8 weeks each year (late May/early June to late July) between 2009 and 2023, where teams repeatedly searched along two valley roads and in the surrounding vegetation and sampled all flowering *Antirrhinum* plants found. The same location was visited 4 to 6 times each year (~every 10-12 days), ensuring we captured plants that started flowering at different times during the season. DNA was extracted from dried leaf material and SNP genotyped using the KASP platform (31) at each of six colour loci including *ROSEA* (ros_assembly_543443), *ELUTA* (ref.9), *SULFUREA* (s91_39699) and *FLAVIA* (s316_93292 and s316_257789), as well as the other divergent loci in recently described colour genes (*RUBIA* - s261_720757 and *CREMOSA* - s1187_290152). The *SULFUREA* marker was chosen as that showing the steepest cline. The frequency of the *SULF^p^* allele did not approach zero on the *striatum* (left) flank most likely because of recombinant alleles (genotyping revealed that 24% of plants judged to be homozygous *sulf^s^* based on phenotype carried the *SULF^p^* marker). The loci were genotyped for a total of >12,000 individuals across 15 years for a larger pedigree study along the hybrid zone and spanning the transition in colour phenotypes from yellow *A.m.m.* var. *striatum* to magenta *A.m.m.* var. *pseudomajus*.

**Cline fitting with simulated annealing**

The hybrid zone at Planoles lends itself to one-dimensional cline fitting due to the narrowness of the habitat along a primarily East – West valley with high mountains to the north and south that are above the altitude limit of these species. Here, *Antirrhinum* plants are primarily found within 100m either side of two roughly parallel roads that run up the valley. First, allele frequencies and genotype counts were determined within a 200 x 200 metre grid of discrete demes. This scale minimised deficits from Hardy-Weinberg equilibrium (see Results) whilst ensuring sufficient sample size within demes (mean = 40, sd = 20, range 10-200) and yielded 125 demes across the hybrid zone. Only one marker within *ROS1* displayed significant departures from HW with a deficit of heterozygotes (*F*>0) even at small spatial scales (<50 metres). Therefore, we expect, based on theory, that this feature may generate narrower cline widths than expected. A more detailed examination of the effects of deme size and departures from HW is presented elsewhere (Surendranadh *et al*., in prep).

The clines were characterised for each colour locus, using a modified version of a custom R script (slowClines) as described (9, 14). This script uses fits a symmetric sigmoid cline with five parameters:

$$\hat{p}=\frac{p_{0}+\left( p_{1}-p_{0} \right)}{1+\exp\left( -4\left( \frac{x-c}{w} \right) \right)}$$

Where $c$ = cline centre, $w$ = cline width (1/gradient), $p_{0}$ = allele frequency at the asymptote in the west (*A.m.m.* var. *striatum* parental allele frequency) and $p_{1}$ = allele frequency at the asymptote in the east (*A.m.m.* var. *pseudomajus* parental allele frequency) and $F_{ST}=var(p)/\bar{p}\left( 1-\bar{p} \right)$. For the $F_{ST}$ parameter, we fitted a beta-binomial error term to account for the variance in allele frequencies across demes and to control for population structure along the cline. Prior to commencing cline fitting algorithm, the larger data set was randomly thinned within each deme to reduce the computational time, resulting in 2900-3300 individual genotypes across ~100 demes.

We use a metropolis-hastings (simulated annealing) algorithm to sample the likelihood surface of the cline fit. We begin the algorithm with a random set of parameters which are changed randomly and the log likelihood *logL* is computed at each iteration. When the next iteration *logL''* has a greater log likelihood than the previous likelihood *logL'* (i.e. *logL''* > *logL'*), the new parameters are accepted. If the next iteration is lower (*logL''* < *logL'*), we accept with a probability *logL''*/*logL'*. To ensure ample exploration of the likelihood surface, the jump size for the next set of parameters are adjusted by a factor of 1.05 when accepted (accept scale) and by (1/1.05) when parameters are rejected (reject scale). After some tests of different accept and rejection scales, we found these values achieved efficient mixing and exploration of the likelihood surface with an acceptance rate ~0.5. This algorithm was run for 50,000 iterations with a burn-in = 2000. From this we find the most likely cline parameters, maximum $\log L$ and assume the likelihood surface follows a chi-square distribution to find -2 $\log L$ max 95% credible regions. We visually inspected the joint likelihood surface for each run. Each run was repeated with randomly chosen starting parameters to ensure reproducibility. More detailed cline fitting of alternative shapes and asymmetries combined with estimates of selection are provided elsewhere (Surendranadh *et al*., in prep).

Prior to cline fitting, the allele frequencies in demes were collapsed to one-dimension, by using a linear transect through the approximate cline centre (*p* = 0.5 isocline of *ROS1*). To search for the optimal transect gradient we compared cline width and maximum log Likelihood (*logL*) values with a range of gradients and intercepts centred on the p = 0.5 isocline through the valley (see Fig x for example) at each of the loci.

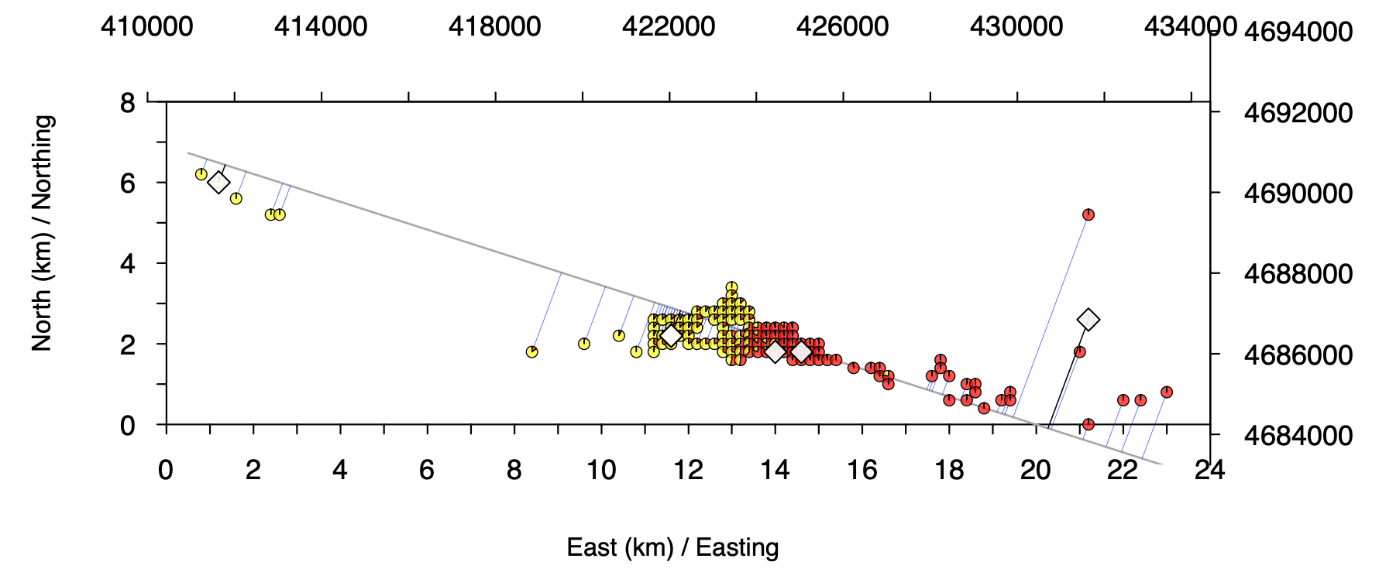

**Fig. a.** Allele frequencies (yellow = *A.m.m.* var. *striatum*, red = *A.m.m.* var. *pseudomajus*) indicated in pie charts at the *ROS1* locus in 200m demes. Each deme is collapsed to a geographic distance along the transect (black line) that is perpendicular (blue line) with the deme. Location of six Whole Genome PoolSeq demes shown as white diamonds. Distance along the hybrid zone show in kilometres and in Easting and Northings. Linear transect show is the optimal line chosen for cline fits with a gradient -0.345 and intercept at 6.9km.

Comparing the optimal transect across loci, there was no common transect direction (gradient) with a best fit across all loci (Fig.b). We found that a gradient of -0.345 and intercept at 6.9km generated the best compromise with highest likelihood (*logL; maxLL*) or within 2 *logL* of the maximum (maxLL) at two of the three loci (*ROS1* and *FLAVIA up*). The third locus (*FLAVIA down*) was a considerably poorer cline fit at this transect, however comparing its best fitting transect cline parameters to the common well-fitting transect (at *ROS1* and *FLAVIA up*) showed similar centres and widths regardless of the transect chosen. Therefore, we report cline parameters for the gradient of -0.345 and intercept at 6.9km.

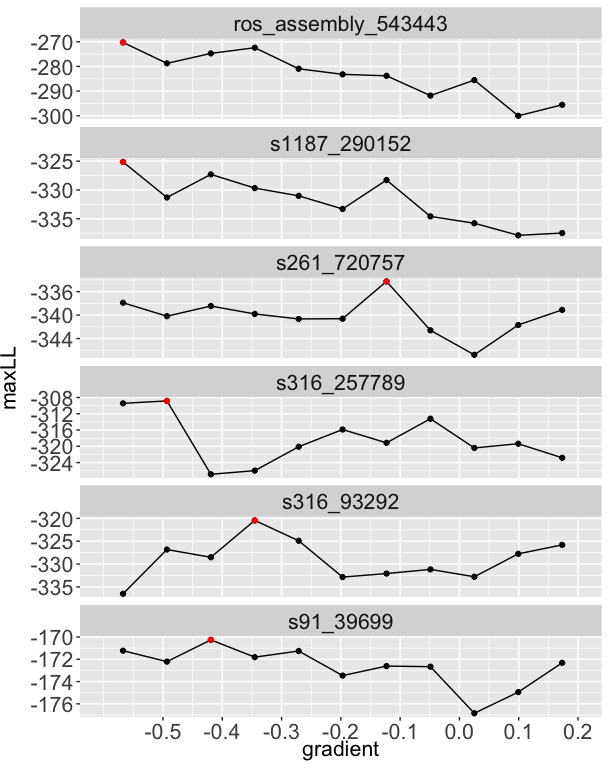

**Fig. b.** Maximum log Liklihood (maxLL) for each parameter search at each transect directions (as gradients) through the hybrid zone at SNP markers at each of six key flower colour loci. The highest maxLL indicated with red filled circle.

**Recombinant haplotype frequencies across the hybrid zone**

At the *FLAVIA* locus we used SNP genotypes flanking either side of the promoter region to distinguish parental haplotypes from the recombinant haplotype. We denote the purebred *A.m.m.* var. *striatum* haplotype as ‘*FLA^s^*’ and the diploid genotype as *FLA^s^/FLA^s^* the *A.m.m.* var. *pseudomajus* as ‘*fla^p^*’ and genotype as *fla^p^/fla^p^*. Recombinant haplotypes are denoted as ‘*FLA^sp^*’. We use two diagnostic SNP markers (s316_93292 and s316_257789), setting genotype scores to reflect the number of copies of the *A.m.m.* var. *pseudomajus* allele, making the three possible genotypes as (i) homozygote genotypes for *A.m.m.* var. *striatum* as 0, (ii) the *A.m.m.* var. *pseudomajus* as 2 and (ii) heterozygotes as 1. Combinations of genotypes at the two diagnostic loci then allowed scoring of parental haplotypes at *FLAVIA* with the following genotype at s316_93292 and s316_257789 as:

- *FLA^s^/FLA^s^* = double *FLA^s^* (parental *A.m.m.* var. *striatum*) = [0,0,]

- *FLA^s^/fla^p^* = heterozygote [1,1]

- *fla^p^/fla^p^* = double *fla^p^* (parental *A.m.m.* var. *pseudomajus*) = [2,2]

- *FLA^sp^/FLA^s^* = *FLA^sp^* recombinant on *A.m.m.* var. *striatum* background = [0,1]

- *FLA^sp^/fla^p^* = *FLA^sp^* recombinant on *A.m.m.* var. *pseudomajus* background = [1,2]

- *FLA^sp^/FLA^sp^* = *FLA^sp^* heterozygous recombinant = [1,1]

This scoring was performed in a custom R script for 2700 individuals and frequencies of haplotypes calculated across the same geographic cline transect as described for diploid genotypes (see above).

**Table.** Geographic cline property estimates from symmetrical sigmoid cline fitting 5-parameter model for six biallelic SNP loci linked to *ROSEA* (ros_assembly_543443), *SULFUREA* (s91_39699) and *FLAVIA* (s316_93292 and s316_257789), as well as the other divergent loci in recently described colour genes (*RUBIA* - s261_720757 and *CREMOSA* - s1187_290152). The best fitting set of parameters are shown, describing the best cline fit with 95% confidence intervals (in parentheses) and include $c$ = cline centre, $w$ = cline width (1/gradient), $p_{0}$ = allele frequency at the asymptote in the west (*A.m.m.* var. *striatum* parental allele frequency) and $p_{1}$ = allele frequency at the asymptote in the east (*A.m.m.* var. *pseudomajus* parental allele frequency). Parameter fits for $F_{ST}=var(p)/\bar{p}\left( 1-\bar{p} \right)$ were conducted but not shown here.

| locus | centre | width | $p_{0}$ | $p_{1}$ |
| --- | --- | --- | --- | --- |
| *CREMOSA* | 13.86 (13.72 – 13.97) | 3.01 (2.73 – 3.43) | 0.05 (0.038 - 0.061) | 0.91 (0.884 - 0.921) |
| *FLAVIA down* | 13.90 (13.78 – 14.10) | 3.00 (2.62 – 3.58) | 0.03 (0.011 - 0.042) | 0.94 (0.913 - 0.967) |
| *FLAVIA up* | 10.57 (10.23 – 10.95) | 9.05 (8.46 – 10.07) | 0.11 (0.089 - 0.184) | 0.99 (0.991 - 0.999) |
| *SULFUREA* | 13.55 (13.55 – 13.56) | 9.34 (9.34 – 12.50) | 0.32 (0.318 - 0.321) | 0.93 (0.929 - 0.939) |
| *RUBIA* | 11.91 (11.46 – 12.43) | 7.57 (5.97 – 9.11) | 0.13 (0.063 - 0.225) | 0.95 (0.898 - 0.968) |
| *ROSEA* | 13.21 (13.14 – 13.26) | 0.88 (0.77 – 1.02) | 0.11 (0.09 - 0.117) | 0.99 (0.976 - 0.991) |

***FLA* recombinant allele characterisation**

To analyse the *FLA* genomic structure and its allelic variants, we selected ~7-8kb either side of the single exon of ~1.5kb at the interval Chr2: 53,460,000-53,476,000. We made WGS of an *A.m.m.* var. *striatum* Accession (Ac1256 MON, Coen ID D125), a *FLA* recombinant line originally from the Hybrid Zone (Coen ID D292) and an *A.m.m.* var. *pseudomajus* Accession (Ac1097 FLO Coen ID D144). The Illumina reads from these lines were mapped against the *A.m. majus* reference genome v4, compared and indels determined by scanning in IGV. Regions of homology were detected by BLAST at NCBI or local BLAST to the reference genome.

To define more precisely the recombination site of the Hybrid Zone *FLA* recombinant allele, revealed by its genomic structure, we used two methods. Firstly, we used PCA results to define a high confidence set of pure homozygotes (n=6 for *pseudomajus* [Accessions with Coen IDs D169, D180, D181, D182, D183, D144] ) and n=7 for *striatum* [Accessions with Coen IDs D125, D186, D187, D151, D153, D155, D157]), with a recombinant line established from the Hybrid Zone with Coen ID D292, for comparison. We called genotypes at biallelic SNVs across the interval 53.275-53.80Mb. Individual samples were required to have a minimum depth of 5 and genotype quality of 15. The resulting 358 SNVs were filtered to select those that were diagnostically different between the two “pure” groups (n=41). The figure below shows a screenshot of the spreadsheet. Each row is a SNP, colour is according to genotype: yellow = HOM REF (*striatum*), magenta= HOM ALT (*pseudomajus*). The pure yellow and magenta columns are the two “diagnostic groupings”, the next is the *FLA* homozygous recombinant. The two SNPs defining this breakpoint were mapped to *A.m. majus* genome v3 at Chr2: 53277698 and Chr2: 53278320, an interval of 622bp.

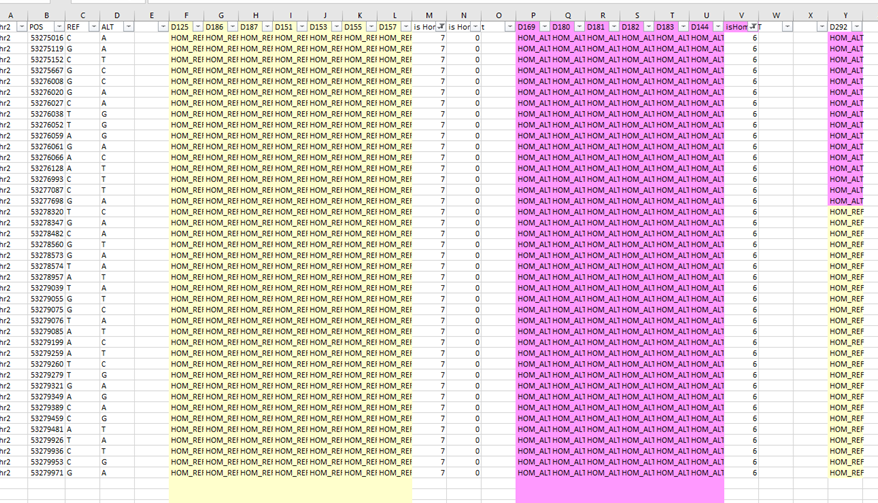

On viewing the region in IGV, we could further refine the interval by identifying an additional diagnostic SNV that had not fit previous filtering criteria. This site had been called as multiallelic since D169 (non-HZ *pseudomajus*) carries the striatum allele here but in phase with additional SNPs that are not seen in *striatum*. This result validated the use of this site to redefine the bounds for v3 as Chr2: 53277698- Chr2: 53277801.

Second, to confirm the above recombination region by Sanger sequencing, we amplified ~2.15kb around the recombination site of various lines using oligos do.259 and do.460 (Supplemental table S1). The lines were: *A.m.m.* var. *pseudomajus* pool MP11, *A.m.m.* var. *pseudomajus* Accessions Ac1110 (CAP: Coen ID D169) and Ac1097 (FLO: Coen ID D144), *A.m.m.* var. *striatum* pool YP4, *A.m.m.* var. *striatum* Accession Ac1256 (MON: Coen ID D125) and allopatric line J1152 (Coen ID D186).

**Supplemental figures. S1 to S8**

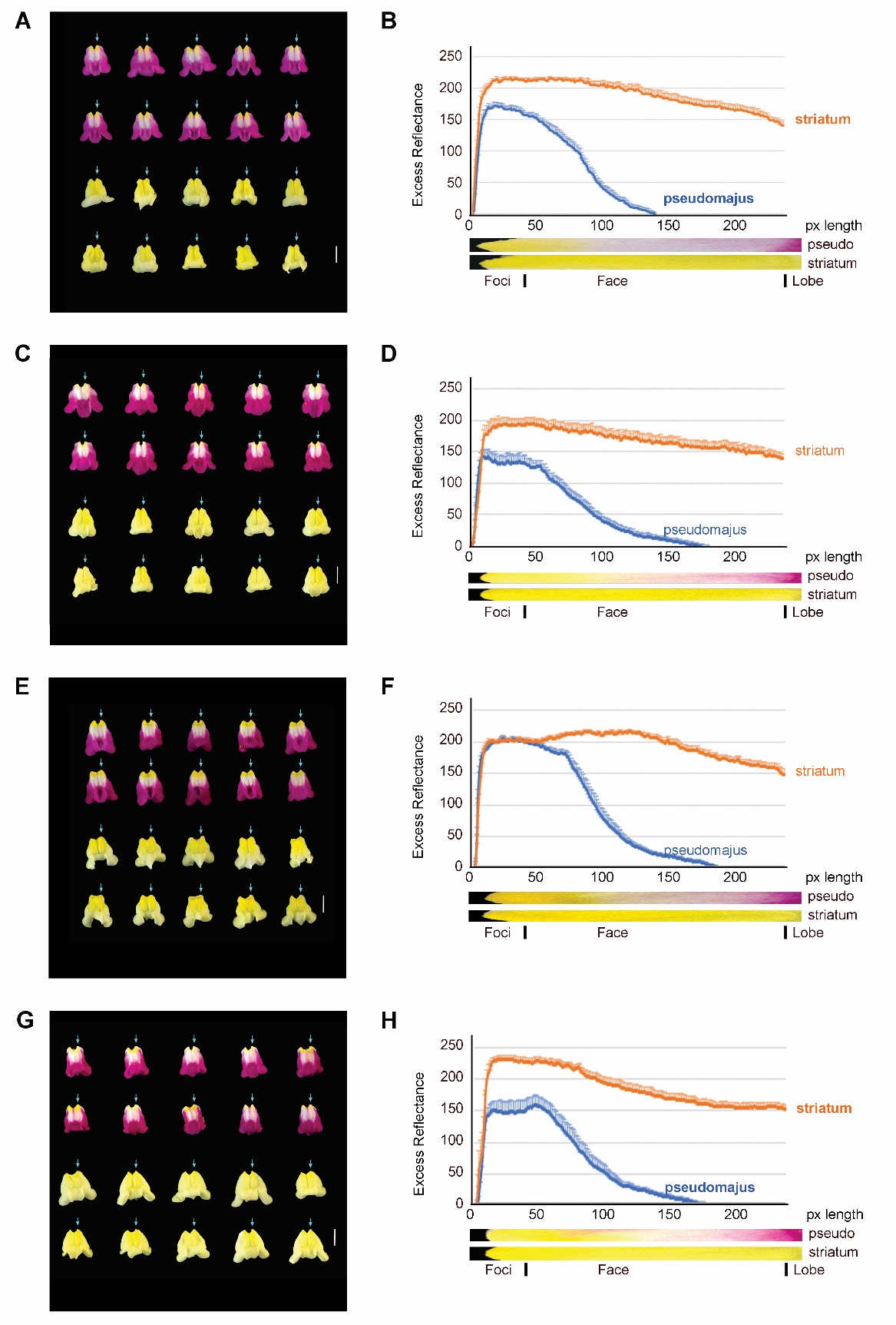

**Supplemental figure S1. Colour Profiles of different *A.m.m.* var. *pseudomajus* and *A.m.m.* var. *striatum* populations.**

**(A)** Seed were collected from wild accessions of *A.m.m.* var. *pseudomajus* (location NDM) and *A.m.m.* var. *striatum* (location VCO) (see location map Supplemental fig.S9). Seed were grown in the greenhouse and, for each accession, two sibs were intercrossed and the progeny sown in 2020. From each population 10 flowers were dissected, photographed and scaled to 1 cm length for foci to the end of lobe (240 pixels in ImageJ). **(B)** The colour profiles of all 10 for each line were determined using the blue arrow as a guide for foci to end of face. Yellow = green reflectance minus blue. Their averages with SE were plotted against distance in pixels from the foci as described (Materials and Methods).

**(C, D)** Analysis as A,B, but using *A.m.m.* var. *pseudomajus* Accession Z-NDM-4 and *A.m.m.* var. *striatum* Z-VCO-6, from different sib intercrosses to those in A,B, and grown in 2023.

**(E, F)** *A.m.m.* var. *pseudomajus* and *A.m.m.* var. *striatum* seed were collected from allopatric populations either side of the Hybrid Zone in the Pyrenees and sown in the greenhouse. Exemplar (archetypal) individuals were selected and sib-intercross seed was generated. After 3 more generations, populations were grown in 2020 and analysed as described in A,B.

**(G, H)** *A.m.m.* var. *pseudomajus* and *A.m.m.* var. *striatum* from the same exemplar parent as E and F, but used different sib crossings and were sown in 2023, with analysis as described in A,B.

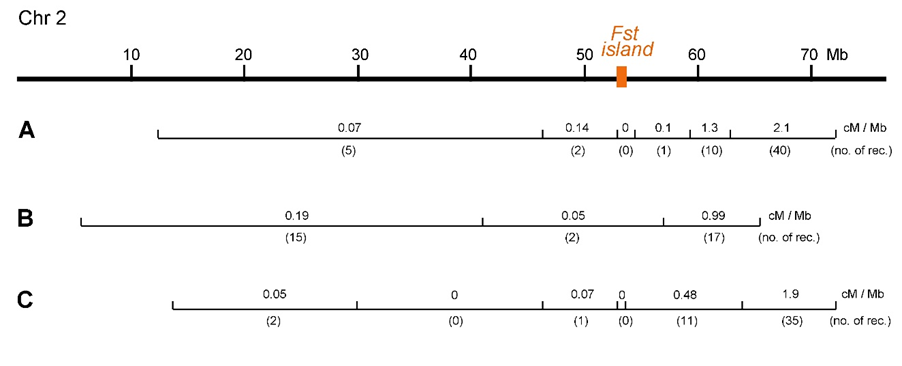

**Supplemental figure S2. Recombination analysis of chromosome 2.** A black horizontal line represents the *Antirrhinum majus majus* reference genome v3 chromosome 2. It is ~76.7 Megabases (Mb) long with an F_ST_ island of ~0.8 Mb (brown rectangle), identified from comparisons between *A.m.m.* var. *pseudomajus* and *A.m.m.* var. *striatum* population genomes. (**A**) From an F2 of *A.m.m.* var. *pseudomajus* x *A.m.m.* var. *striatum* (n=105) we used 10 markers along chr2 to find the number of recombinations for each interval (no. of rec.) to estimate the recombination frequency in centiMorgans/Mb. (**B**) The same analysis was made for an F2 of *A.m.m.* var. *striatum* x *A.m. majus* (n=112). (**C**) The same analysis was made for an F2 of *A.m.m.* var*. pseudomajus* x *A.m. majus* (n=112).

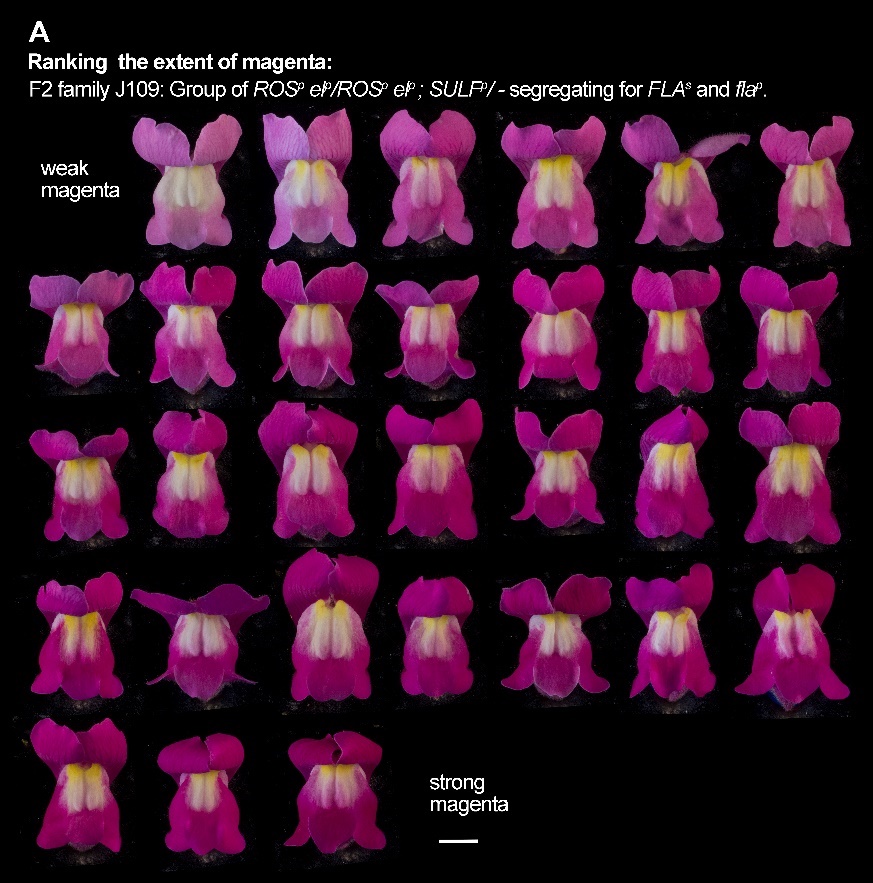

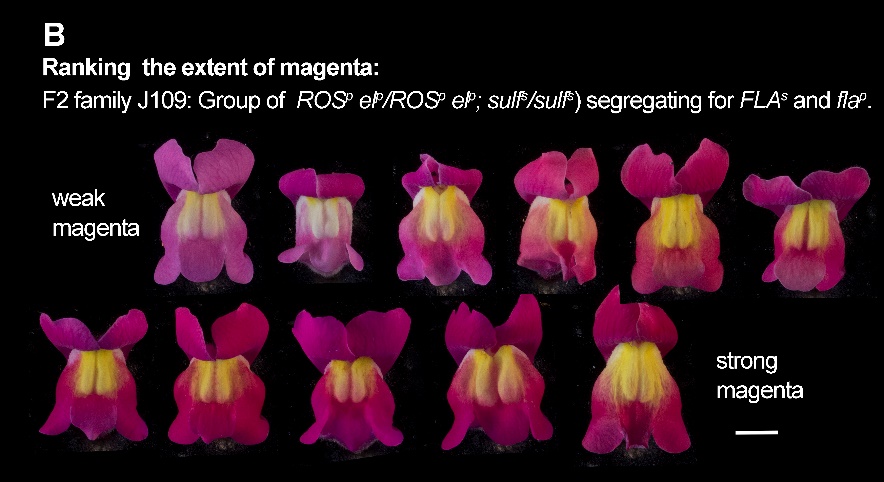

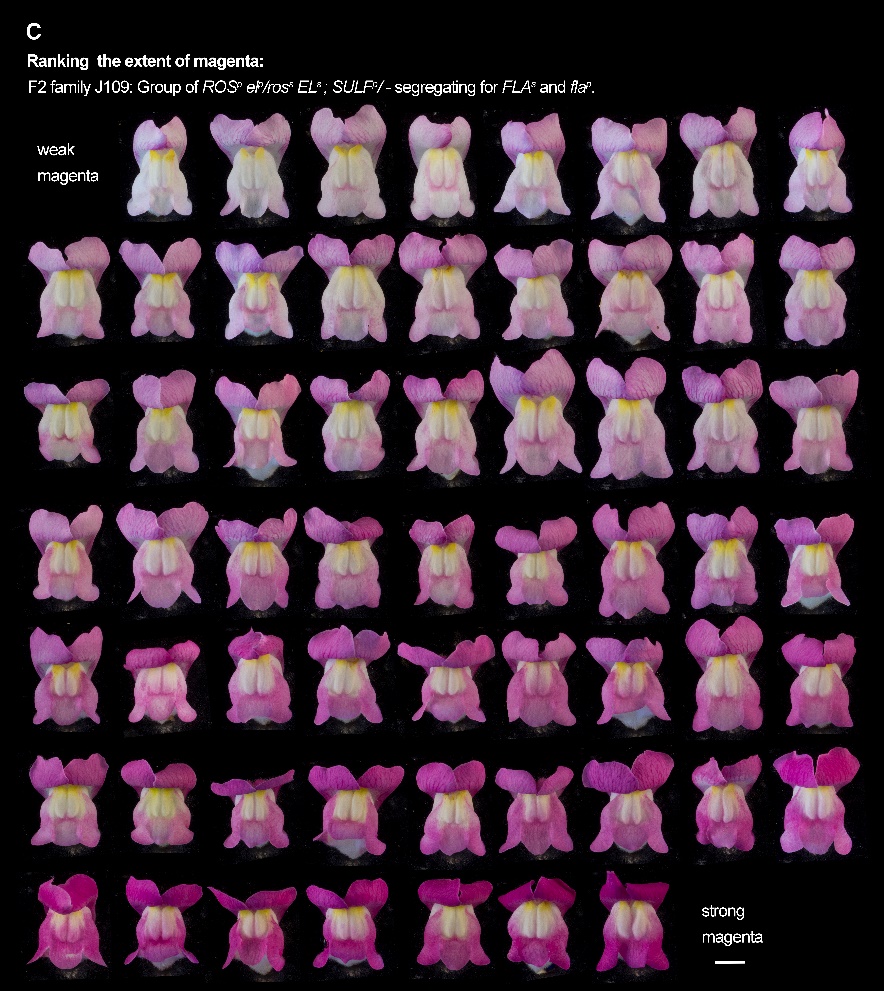

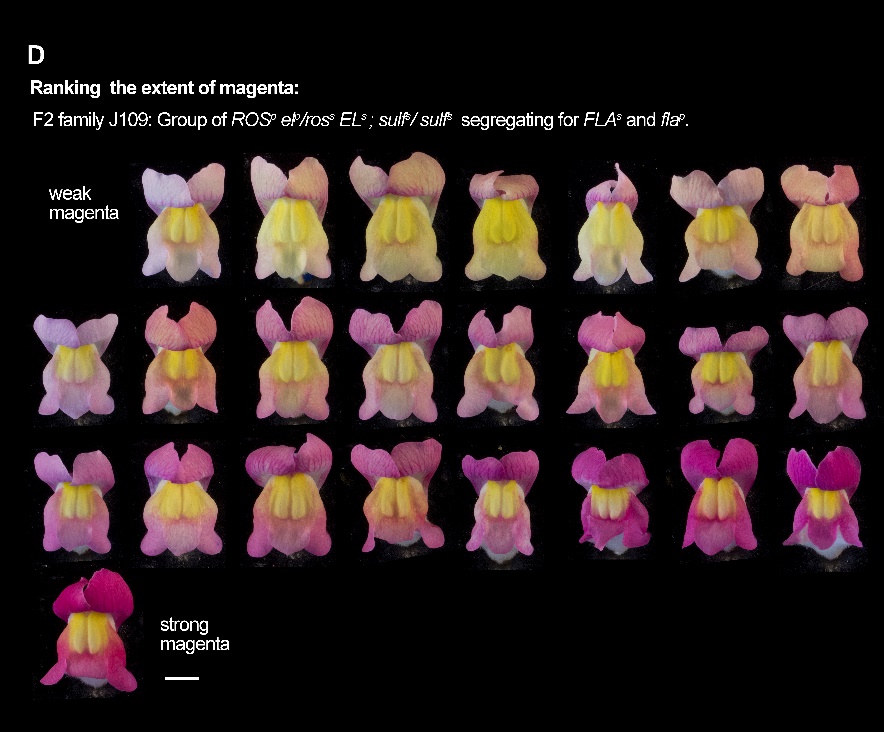

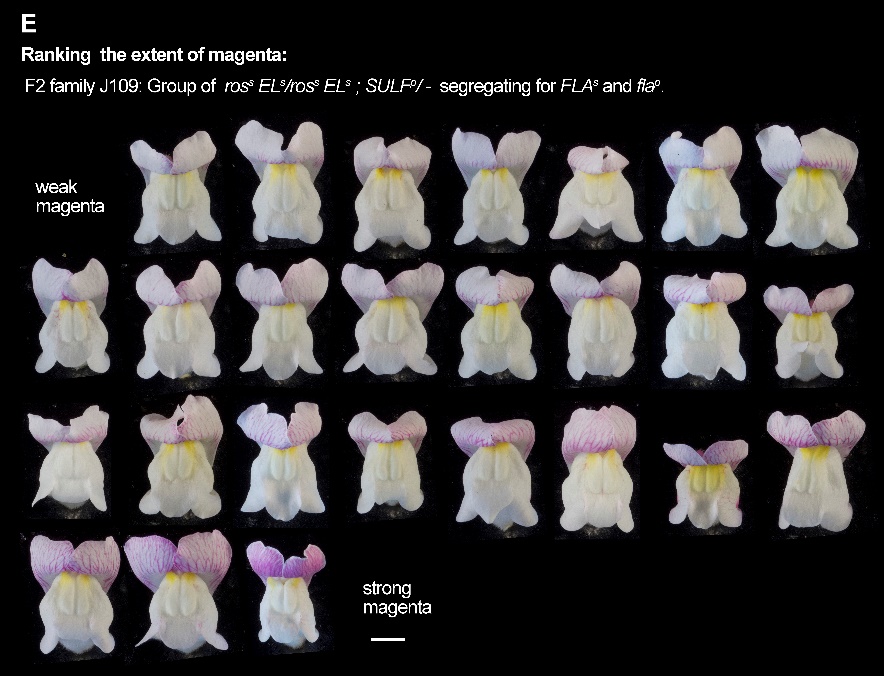

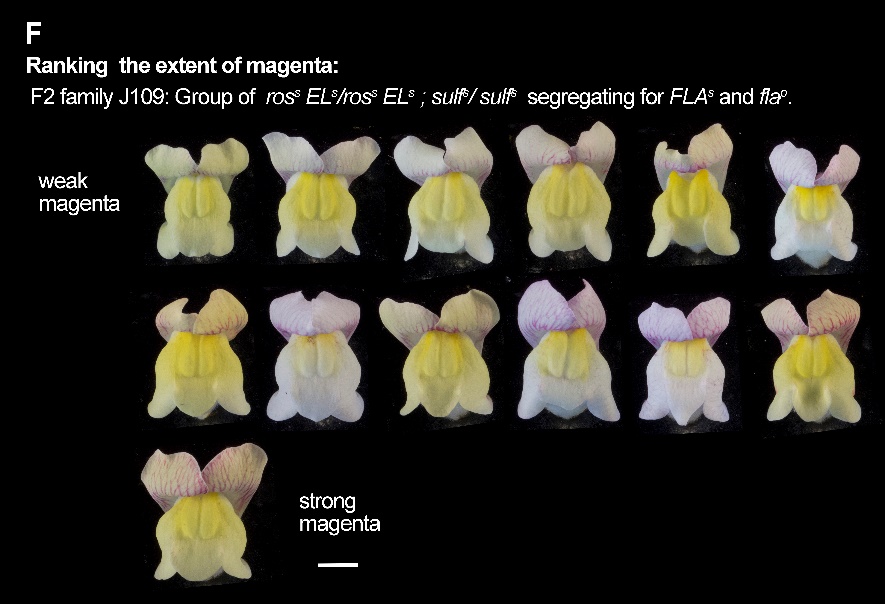

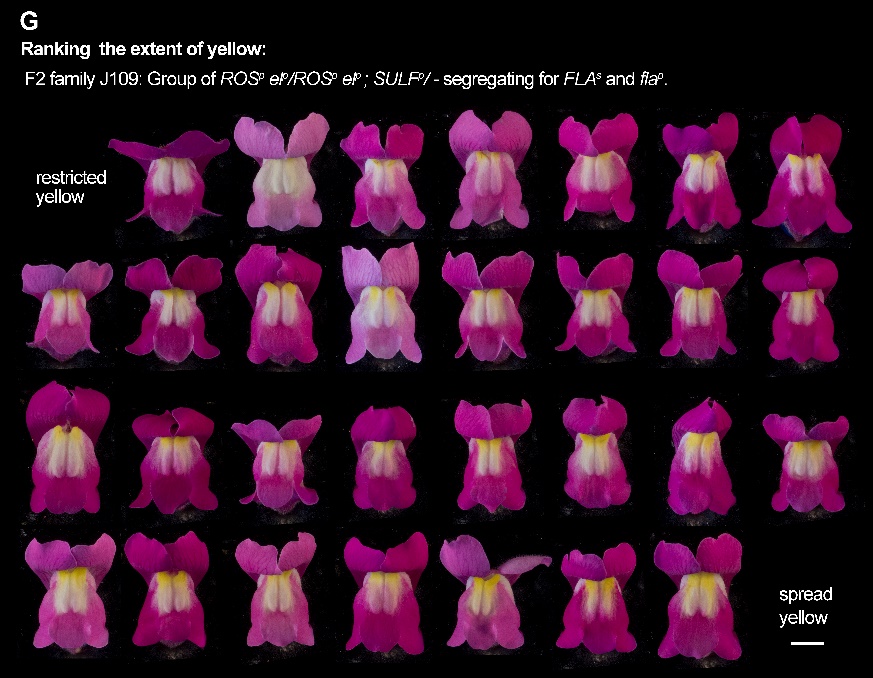

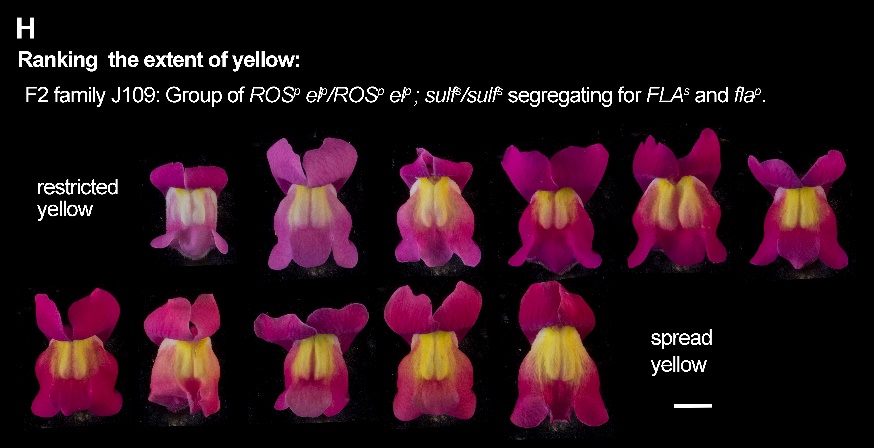

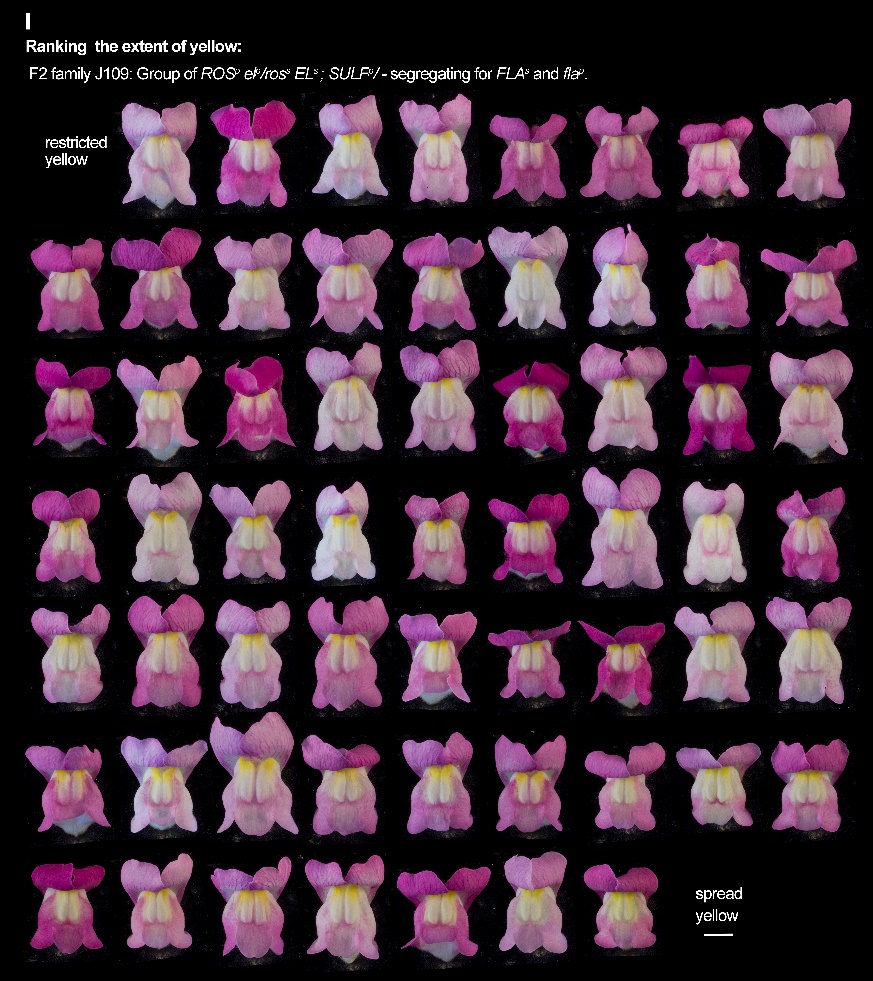

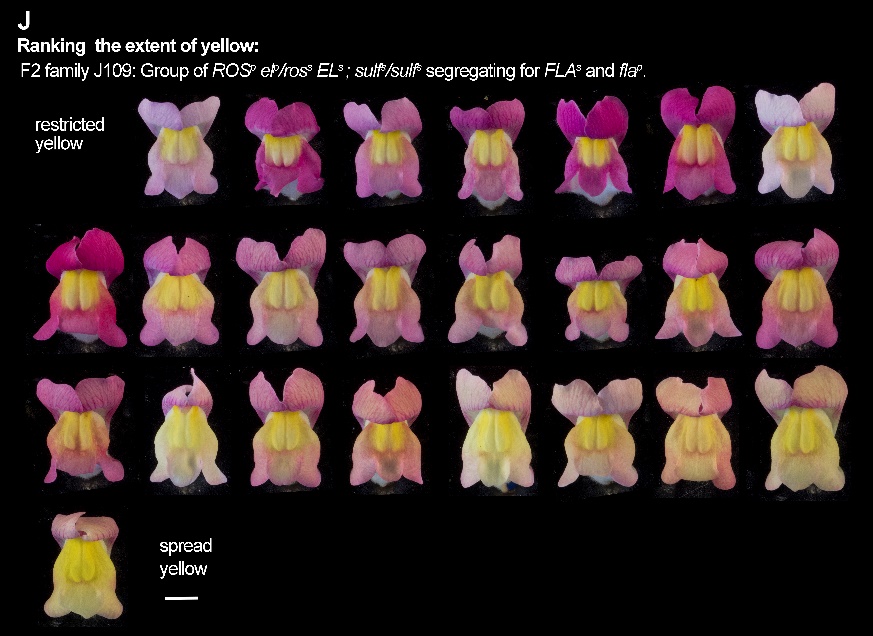

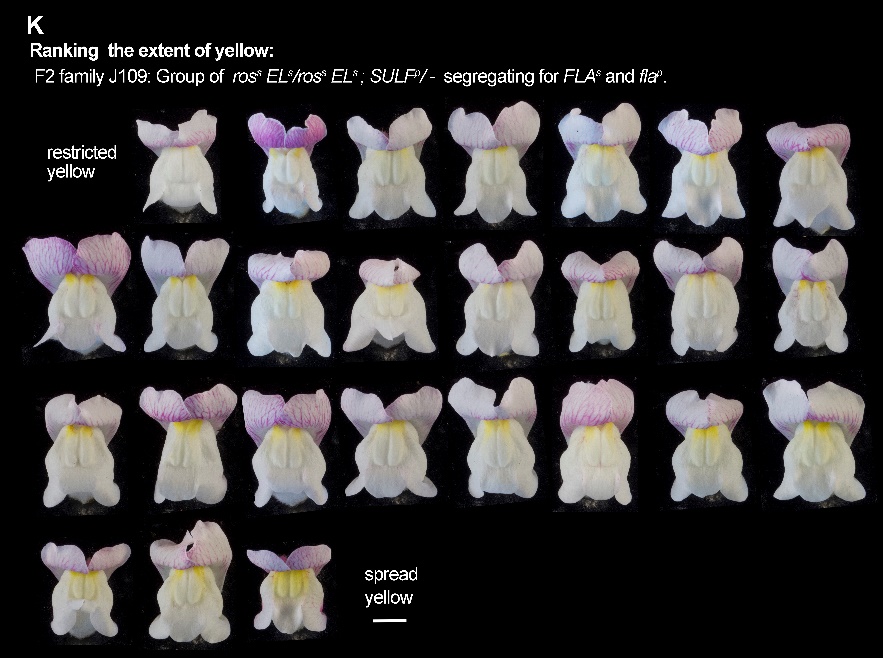

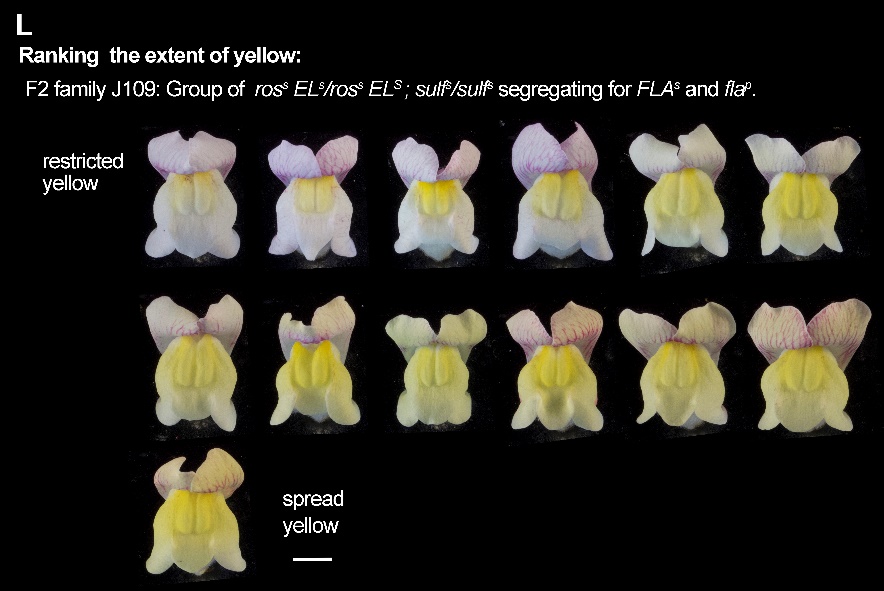

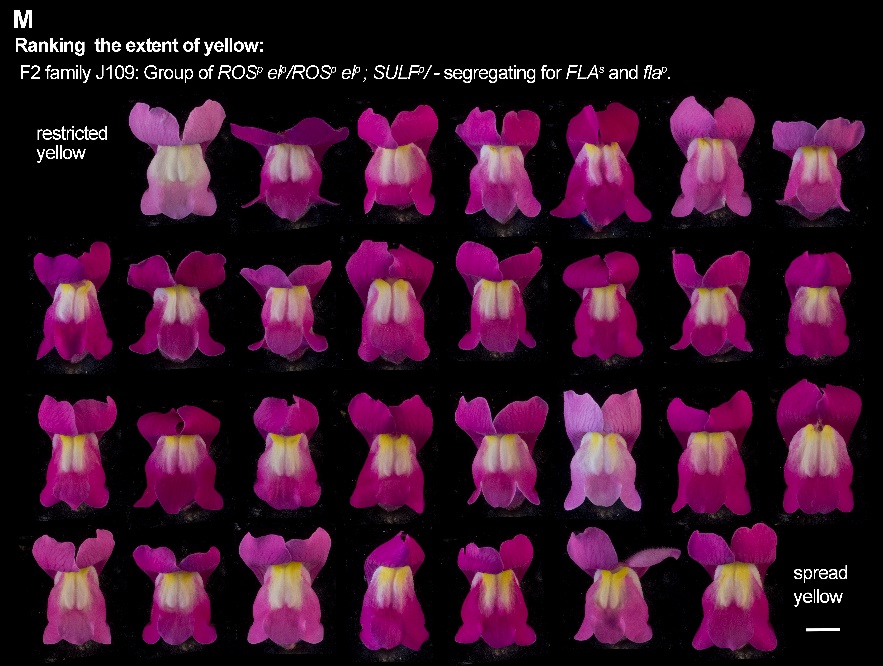

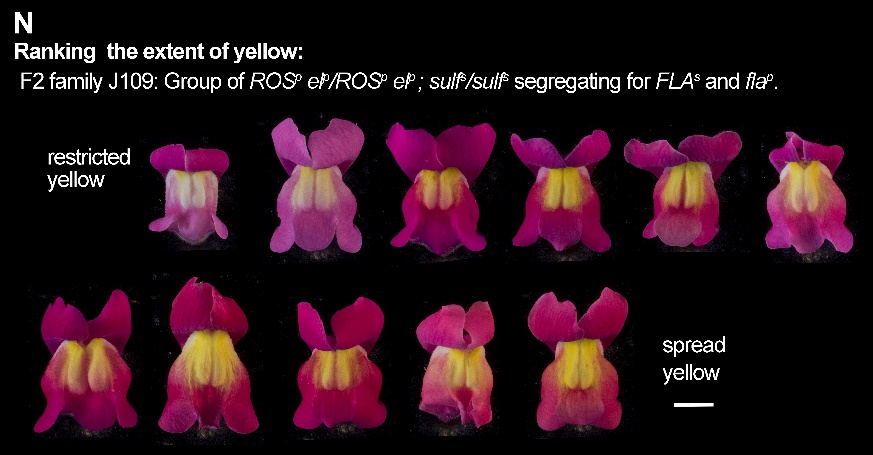

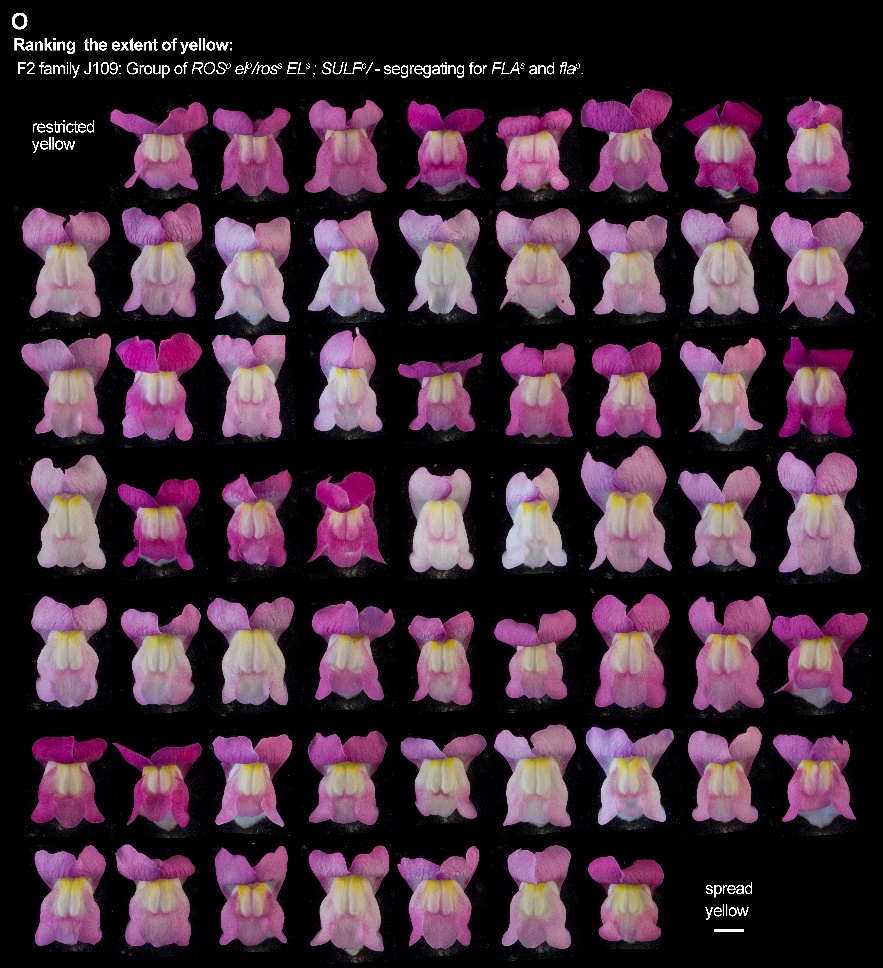

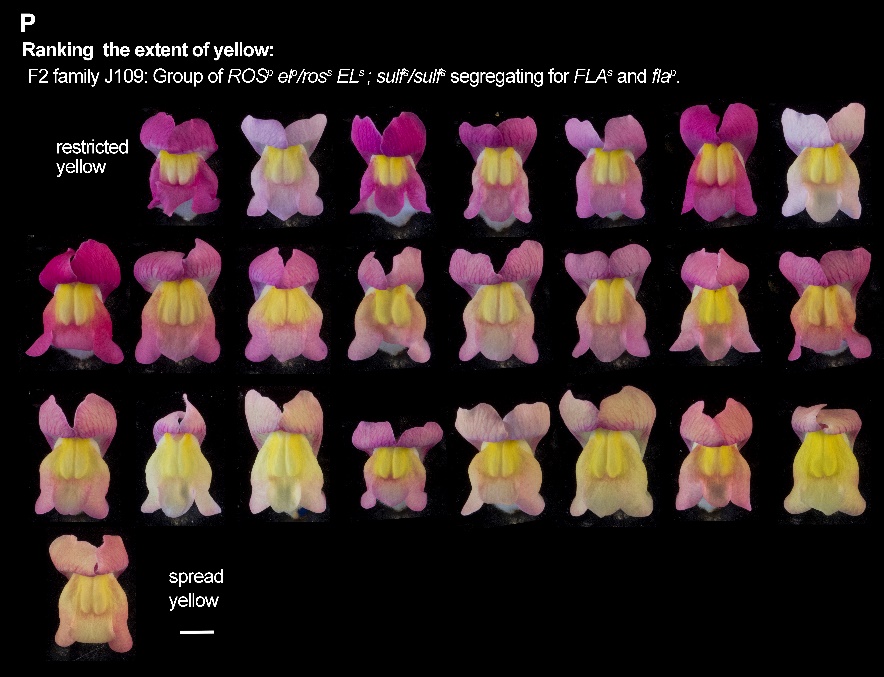

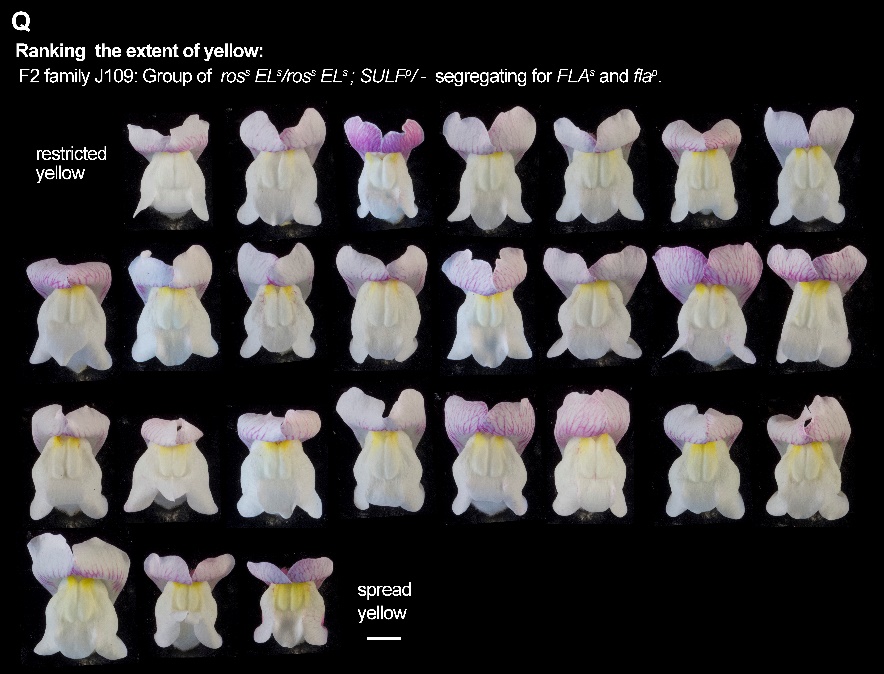

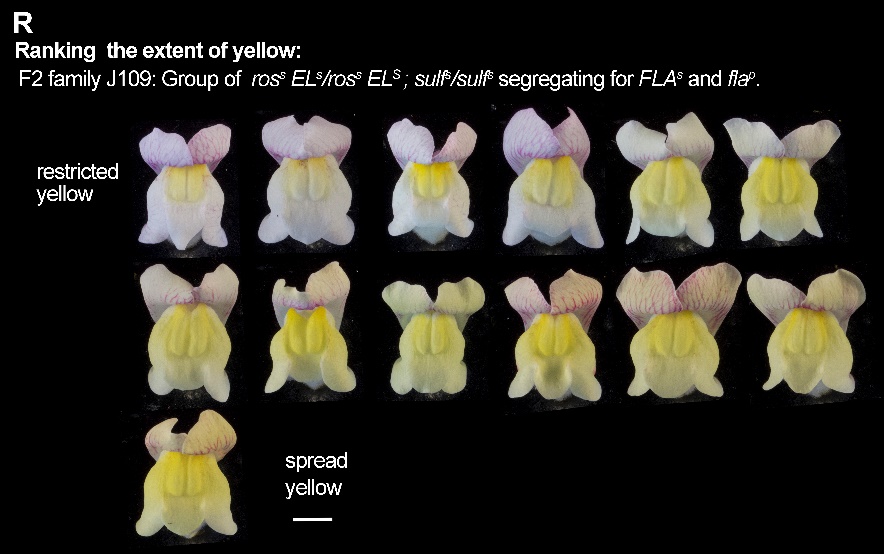

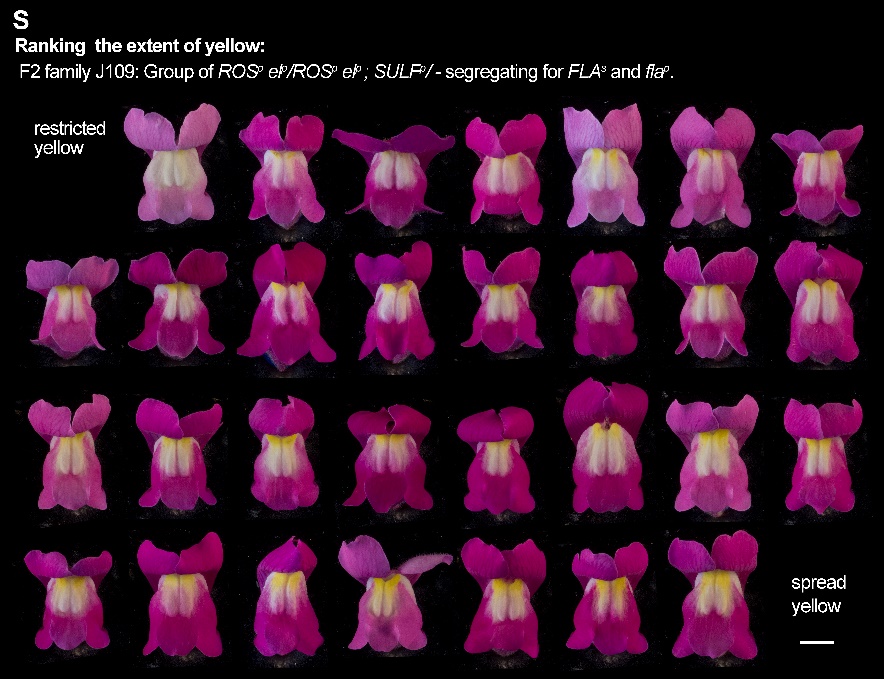

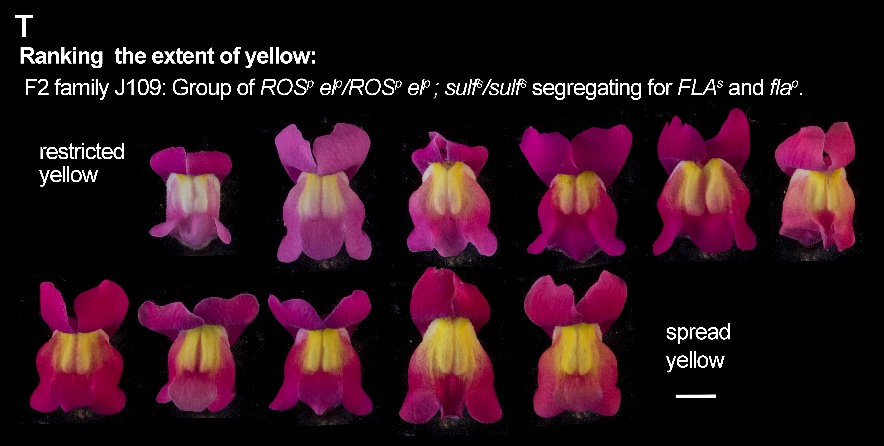

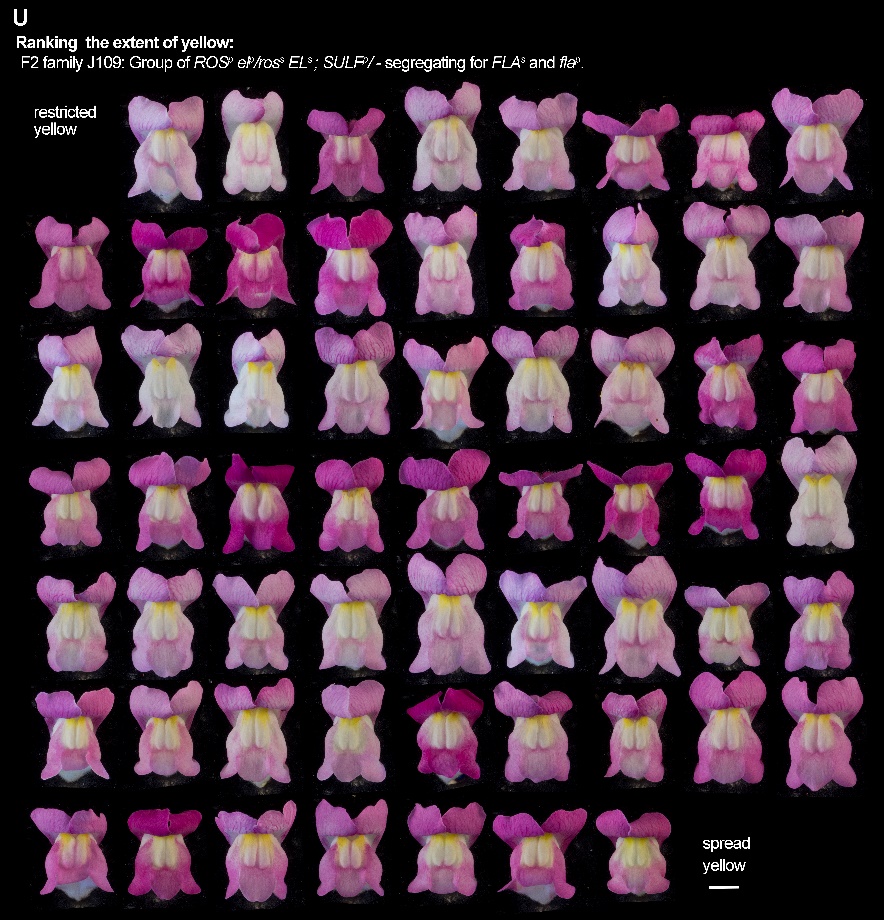

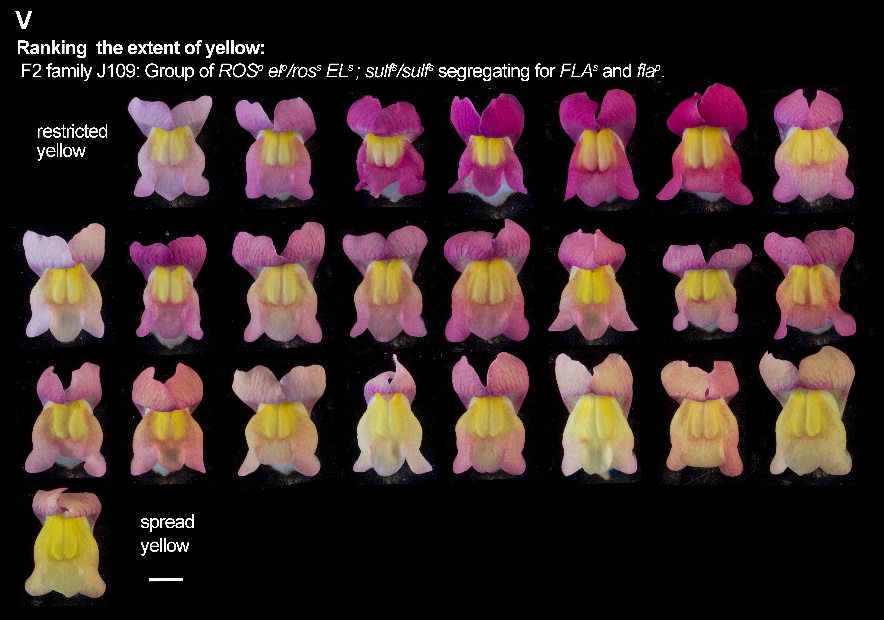

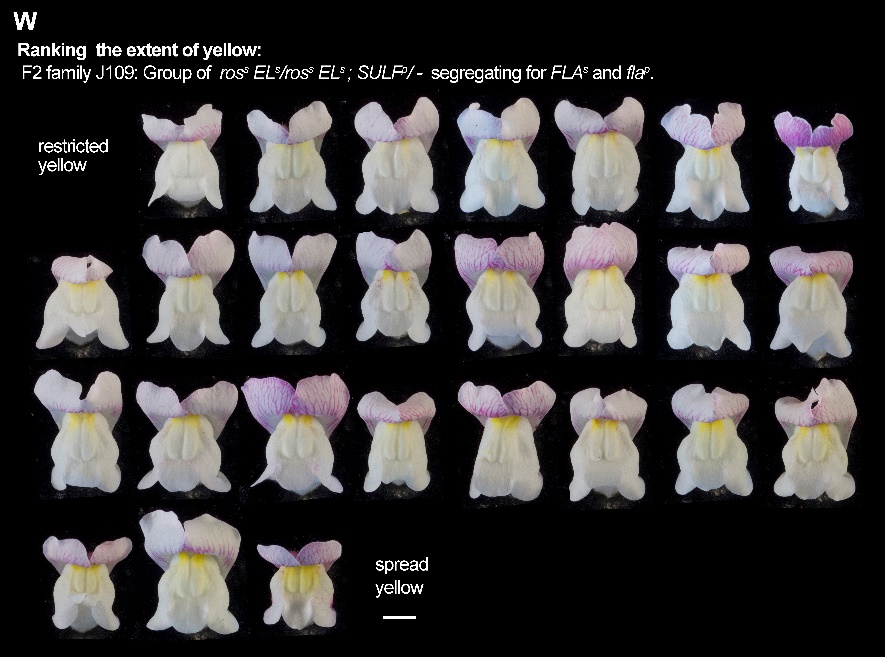

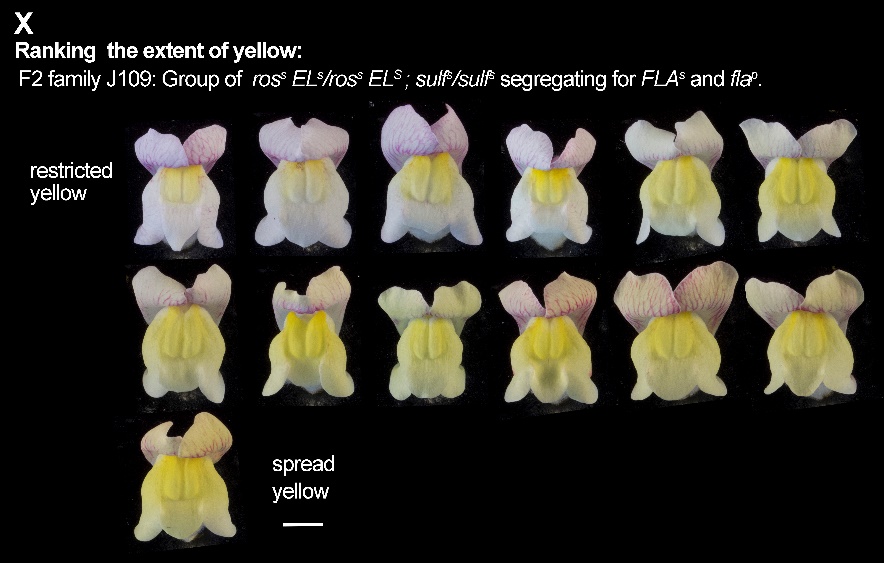

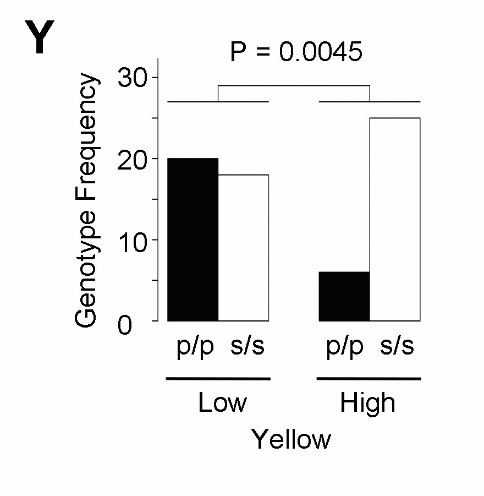

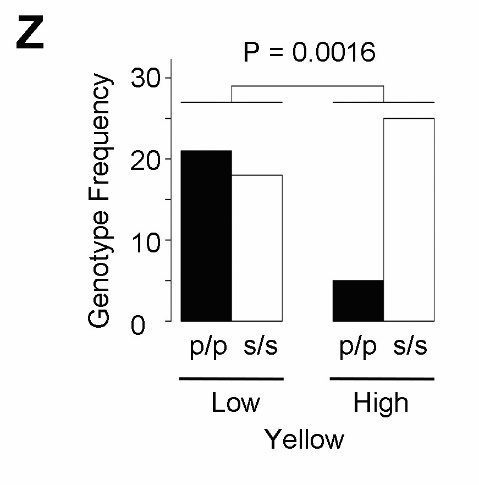

**Supplemental figure S3. Flowers ranked by extent of colour: family J109 (F2 of *A.m.m.* var. *pseudomajus* x *A.m.m.* var. *striatum*).** (**A-F**) Six different genotyped groups, where *FLA^s^* and *fla^p^* were segregating, were separately ranked for the extent of their magenta pigmentation. (**G-L**). The same groups were also ranked for their extent of yellow pigmentation and results analysed as for magenta. In each case, rankings were made without knowing the FLA genotypes. Three researchers, Daniel, Desmond and Tingting, made independent rankings for yellow: Daniel (G-L), Desmond (**M-R**) and Tingting (**S-X**). Similar results were found for all 3 yellow rankings: Daniel data (see Fig.2), Desmond (**Y**) and Tingting (**Z**). Each genotype group was ranked separately and the results then aggregated for all classes. The high yellow phenotypes were significantly enriched for s/s homozygotes with chi-squared P values shown.

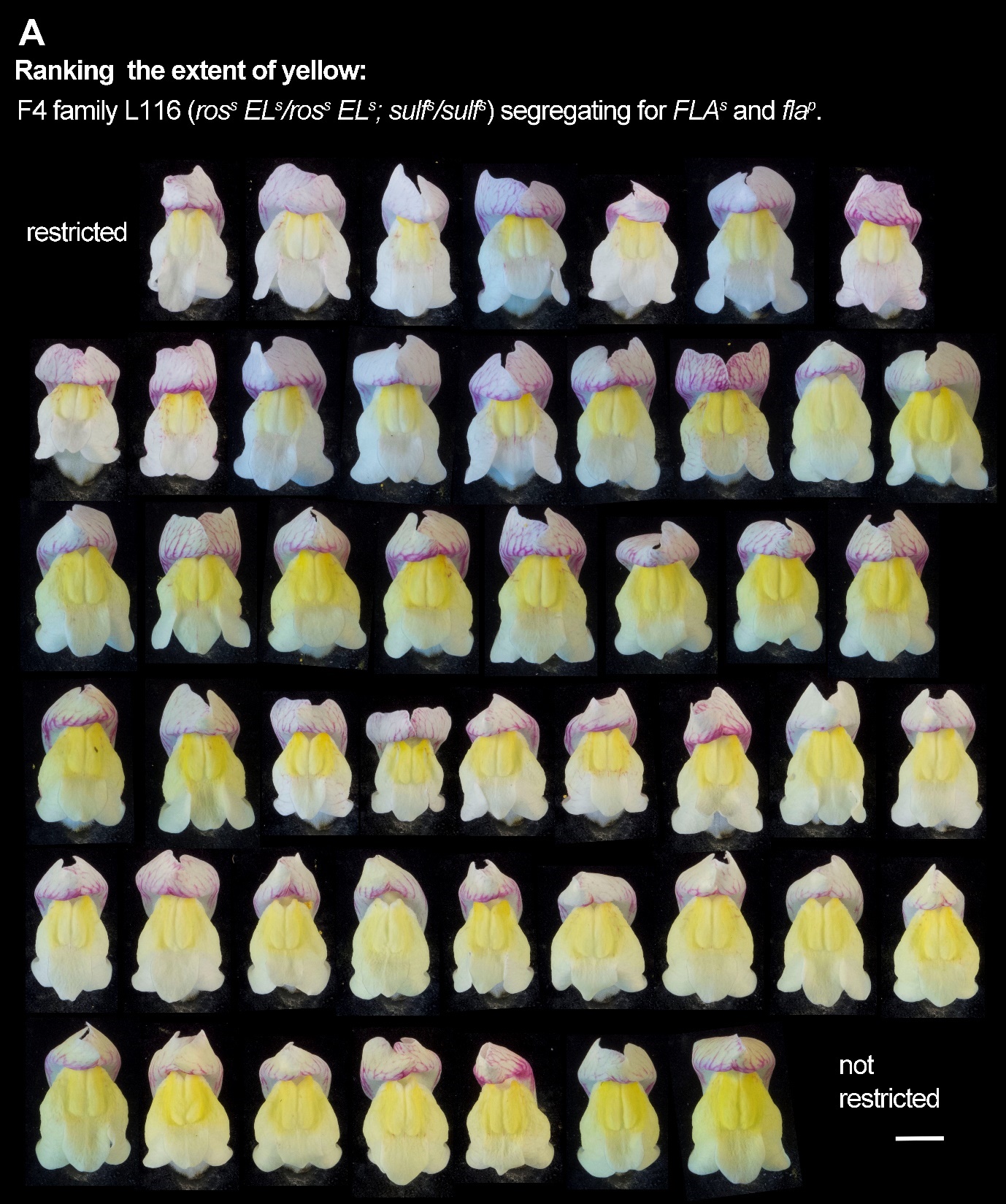

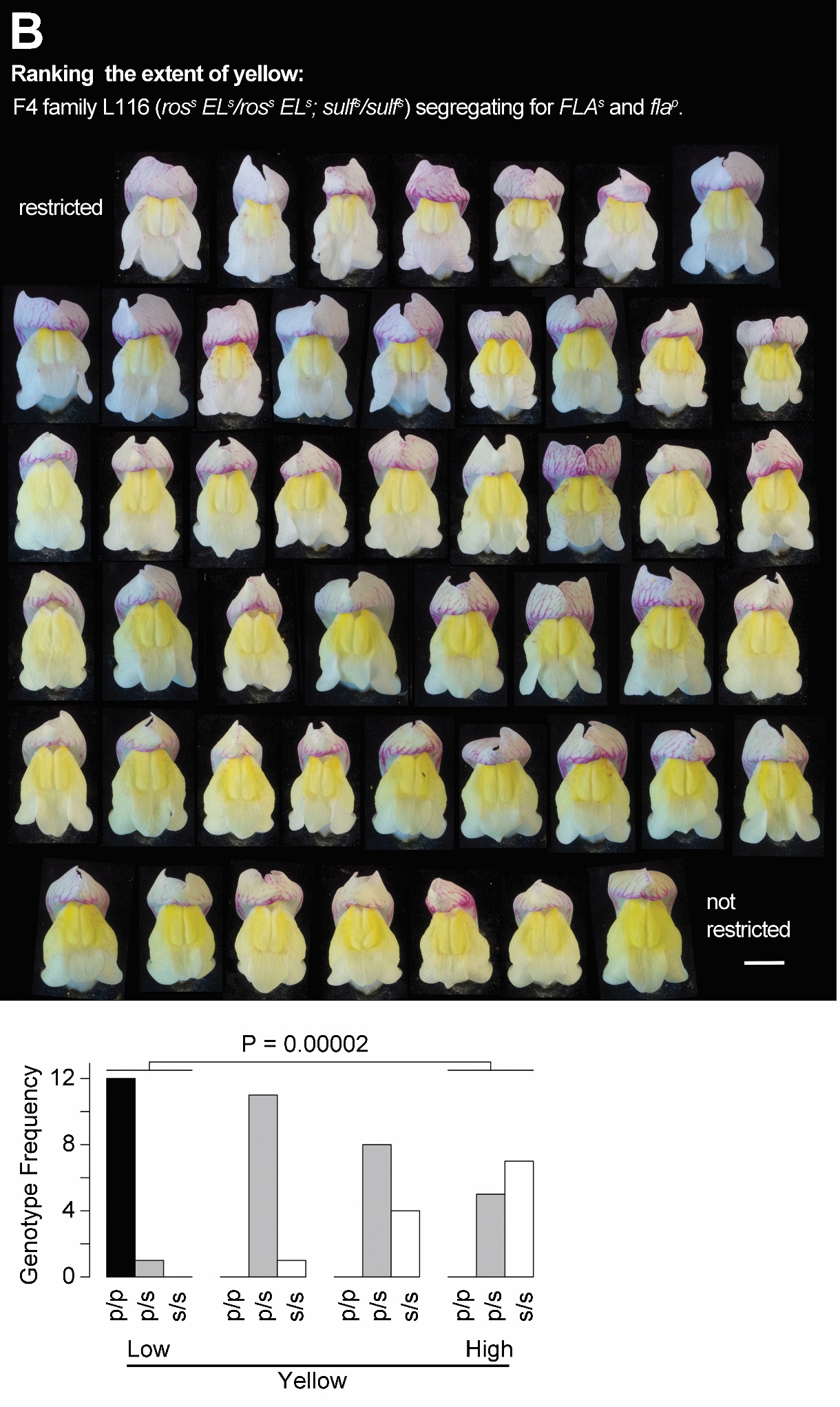

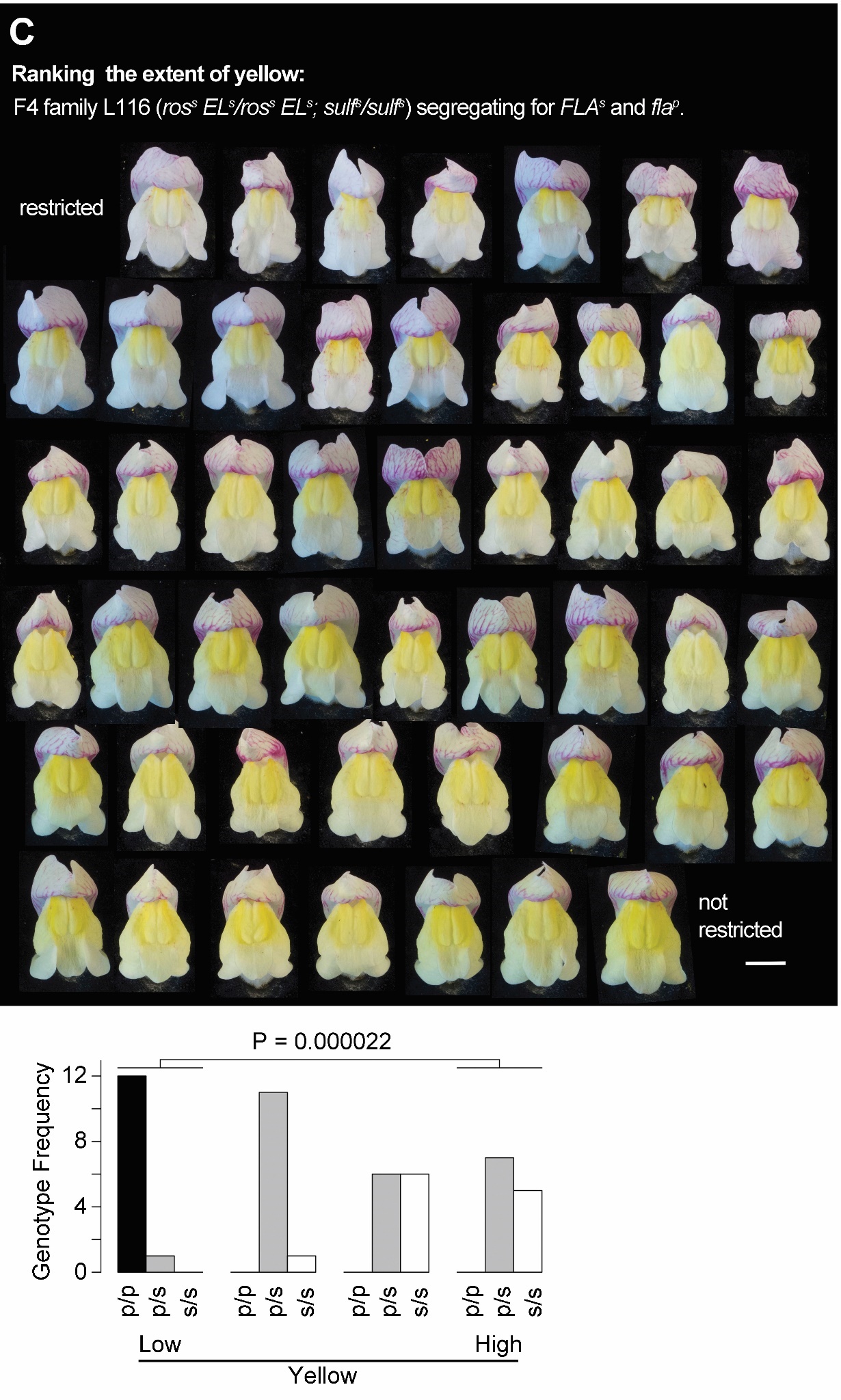

**Supplemental figure S4. Flowers ranked for yellow colour: F4 family L116 (*ros^s^ El^s^/ ros^s^ El^s^ sulf^s^/sulf^s^*).** Family L116 was segregating for *FLA^s^* and *fla^p^*. (**A**) A flower from each individual plant was ranked for its extent of yellow throughout its petal lobes. The ranked plants were grouped into 4 quartiles, their genotypes determined and plotted in Fig.2C. (**B**) Ranked flowers and graphed data from a second person. (**C**) Ranked flowers and graphed data from a third person.

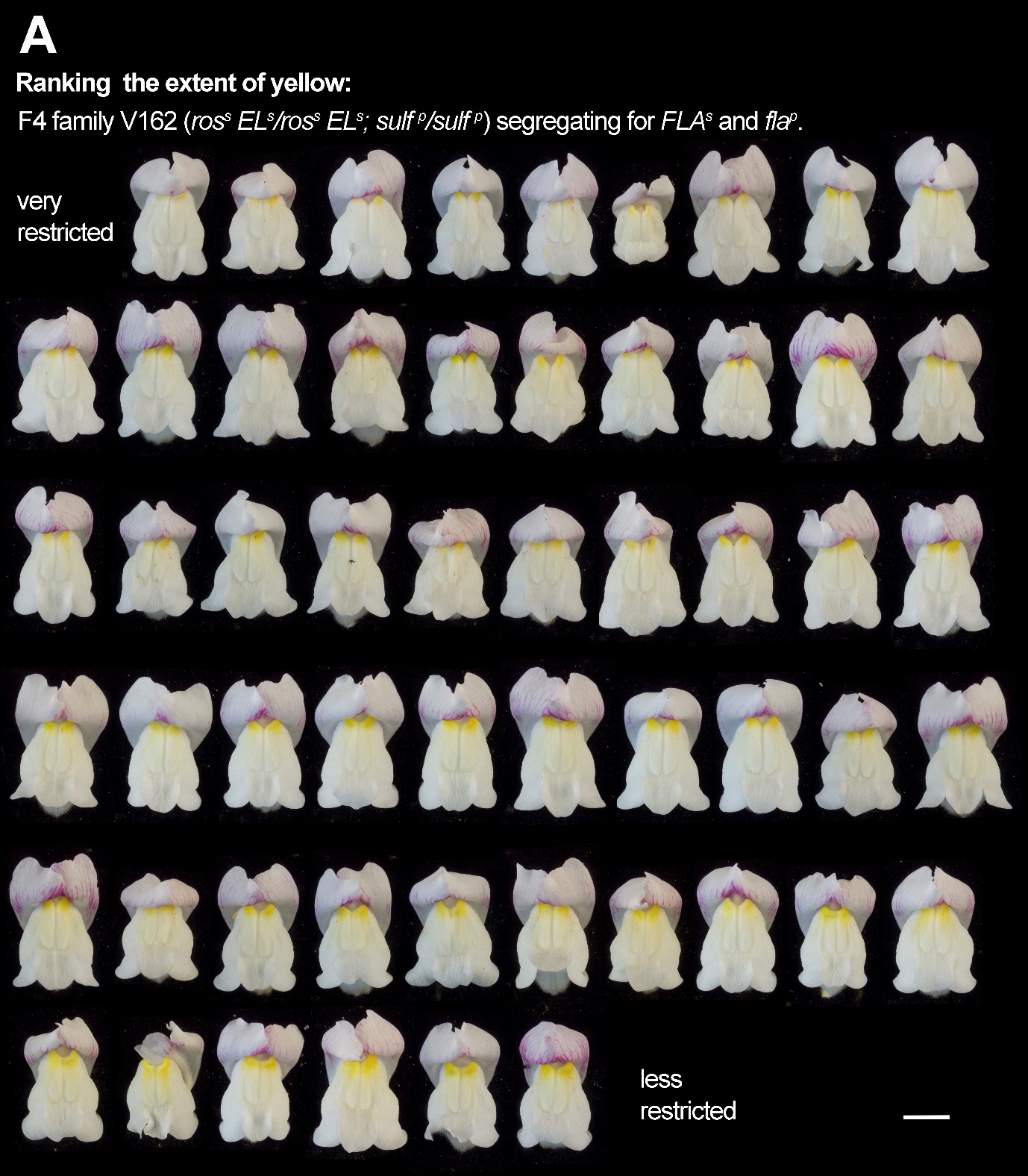

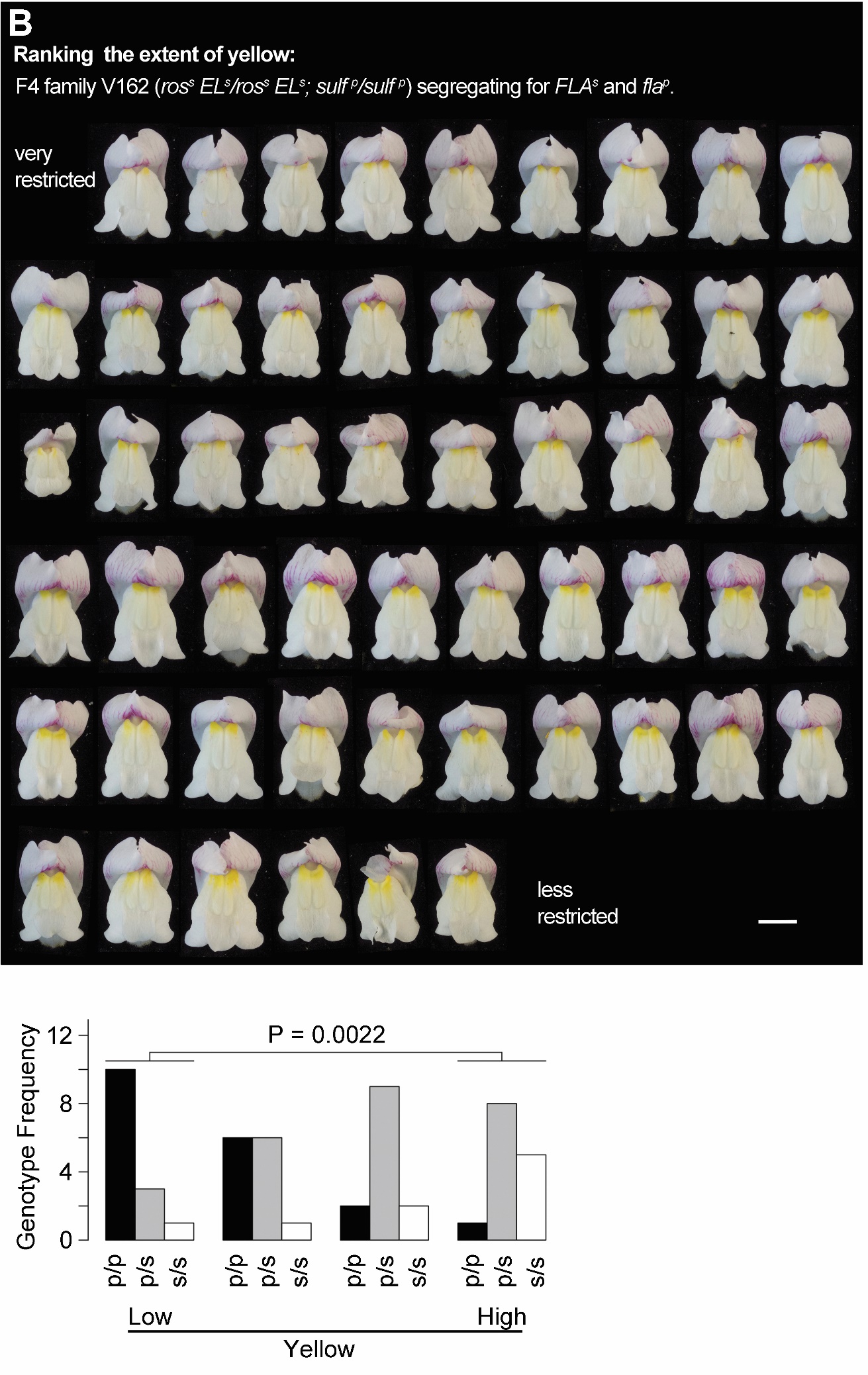

**Supplemental figure S5. Flowers ranked for yellow colour: F4 family V162 (*ros^s^ El^s^/ ros^s^ El^s^ SULF^p^/SULF^p^*).** Family V162 was segregating for *FLA^s^* and *fla^p^*. (**A**) A flower from each individual plant was ranked for its extent of yellow throughout its petal lobes. The ranked plants were grouped into 4 quartiles, their genotypes determined and plotted in Fig.2D. (**B**) A second person’s independent ranking of flowers and graphed data as described in Fig.2D with chi-squared P value determined for lowest versus highest quartiles.

Ranking results from an F2 population (n=69) of A.m.m. var. pseudomajus (p) x A.m.m. var. striatum (s) ranked for extent of petal lobe magenta and genotyped for SNP2A. Individuals were grouped and ranked separately according to ros, el, and sulf genetic backgrounds, and results were aggregated across all genotypic classes. The low and high magenta categories have no significant difference in frequencies of p/p and s/s homozygotes (P values calculated using a contingency chi-squared test between the low and high quartiles). (B) Ranking the same population for extent of yellow shows s/s homozygotes significantly enriched in the high yellow category. (C) A ross Els/ross Els sulfs/sulfs population genotyped for SNP2A and ranked for the extent of yellow. The lowest yellow quartile is significantly enriched for p/p homozygotes. Examples of flowers from the middle of each quartile are shown below. Bar is 1 cm. (D) A ross Els/ross Els SULFp/SULFp population ranked for their extent of yellow. The lowest yellow quartile is significantly enriched for p/p homozygotes. Bar is 1 cm. Yellow rankings in B and C were done in triplicate by 3 independent observers and all gave similar significant P values (fig.S3).

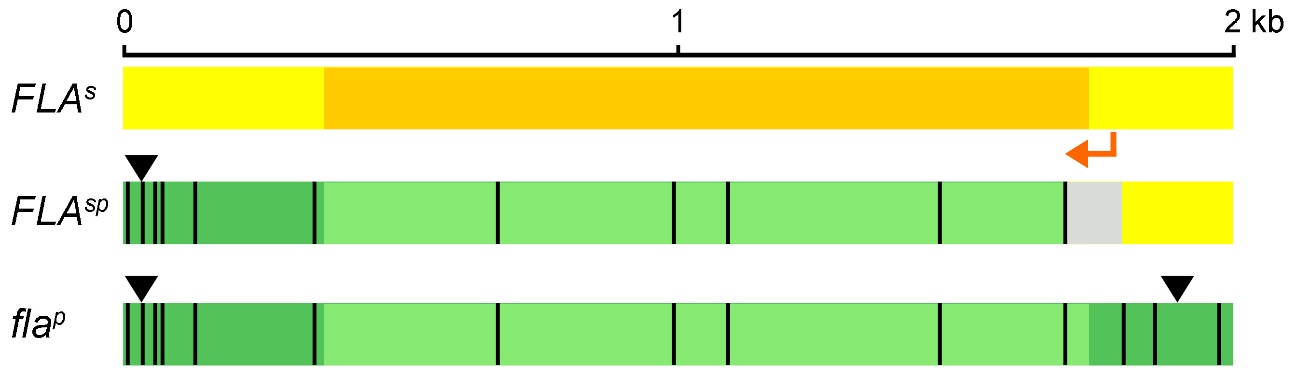

**Supplemental figure S6. Hybrid Zone Recombinant Structure.** The *FLA* region is shown for hybrid zone alleles of *A.m.m.* var. *striatum* (*FLA*^s^), *A.m.m.* var. *pseudomajus* (*fla*^p^) and a recombinant (*FLA^sp^*) found at high frequency in the hybrid zone. The *FLA* coding region (different shade) is *fla*nked by ~300-400 bp of 5’ sequence and 3’ sequence, with SNPs (relative to *FLA*^s^) marked as black lines, and indels as black triangles. The orange arrow indicates transcription direction. The hybrid zone recombination event has joined the 5’ region of *FLA*^s^ to the coding and downstream region of *fla*^p^. As the *FLA^s^* and *fla^p^* sequences are identical near the start region, the recombination point must lie between the two closest SNPs shown by the grey region.

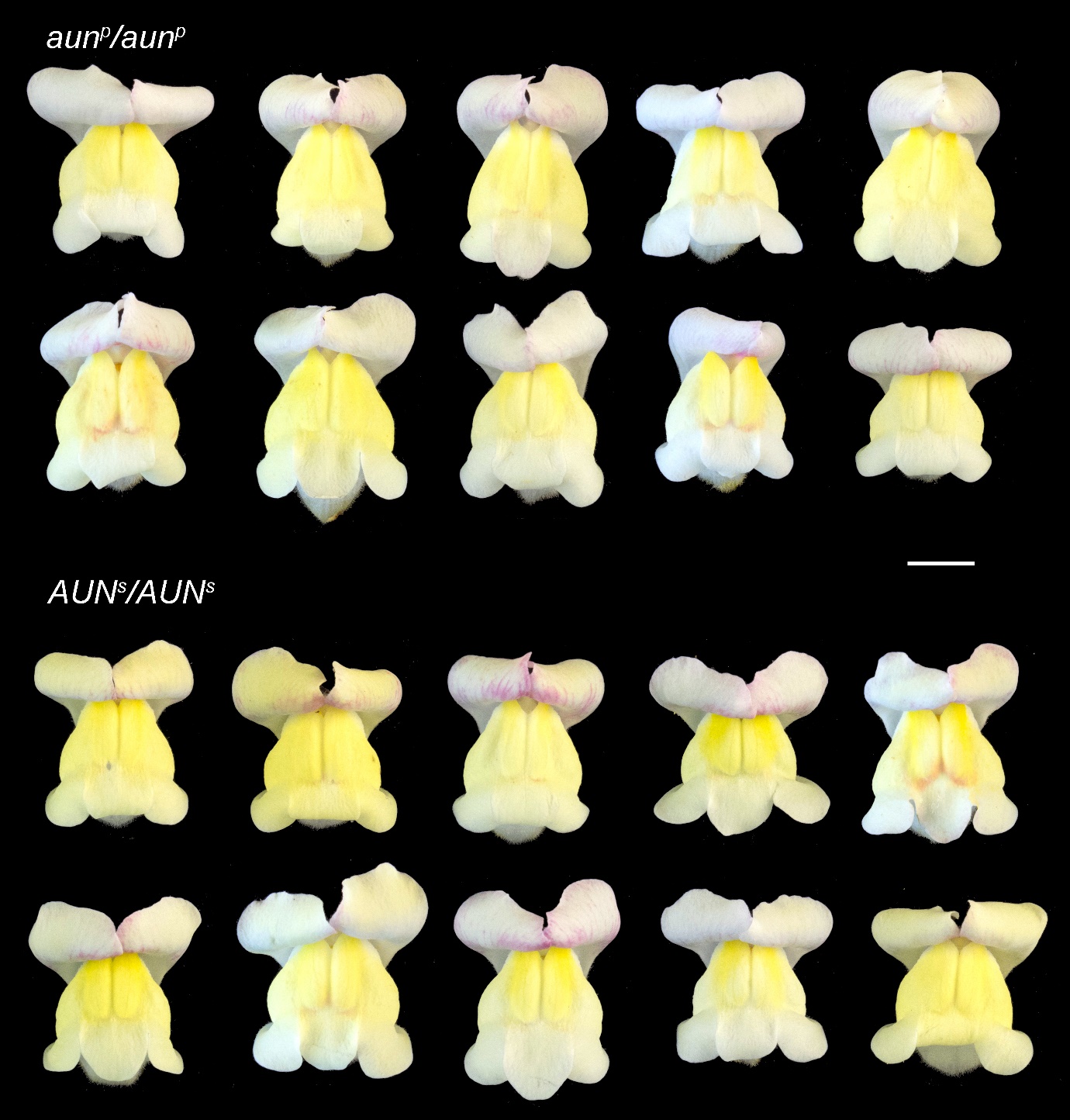

**Supplemental figure S7. Example flower photographs for AUN alleles.** Photographs of *AUN^s^/AUN^s^* and *aun^p^/aun^p^* from 10 different plants in a *sulf^s^/sulf^s^ ros^s^/ros^s^ FLA^s^/FLA^s^ CRE^p^/CRE^p^* background.

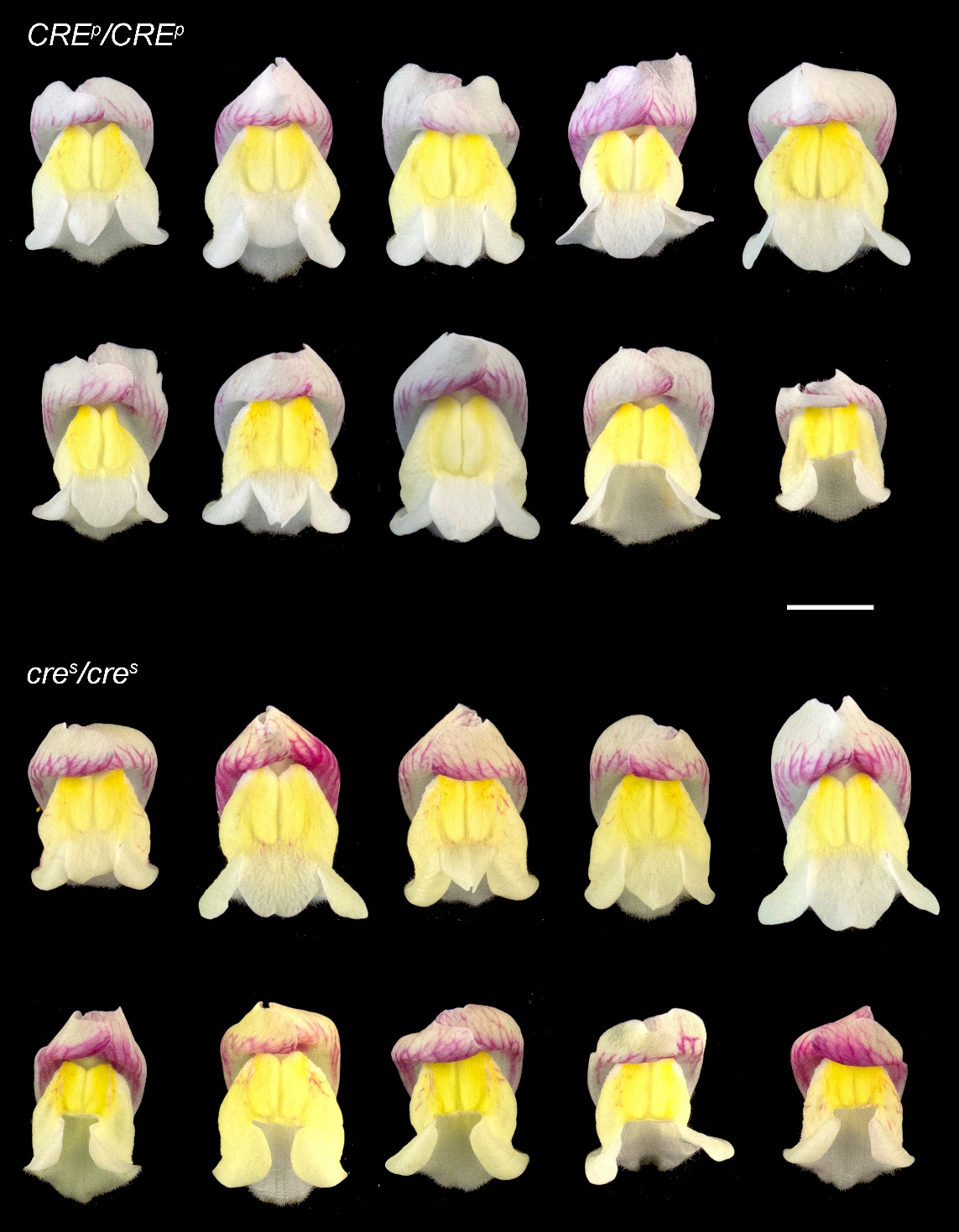

**Supplemental figure S8. Example flower photographs for CRE alleles.** Photographs of *CRE^p^/CRE^p^* and *cre^s^/cre^s^* from 10 different plants in a *sulf^s^/sulf^s^ ros^s^/ros^s^ FLA^s^/fla^p^ AUN^s^/aun^p^* background.

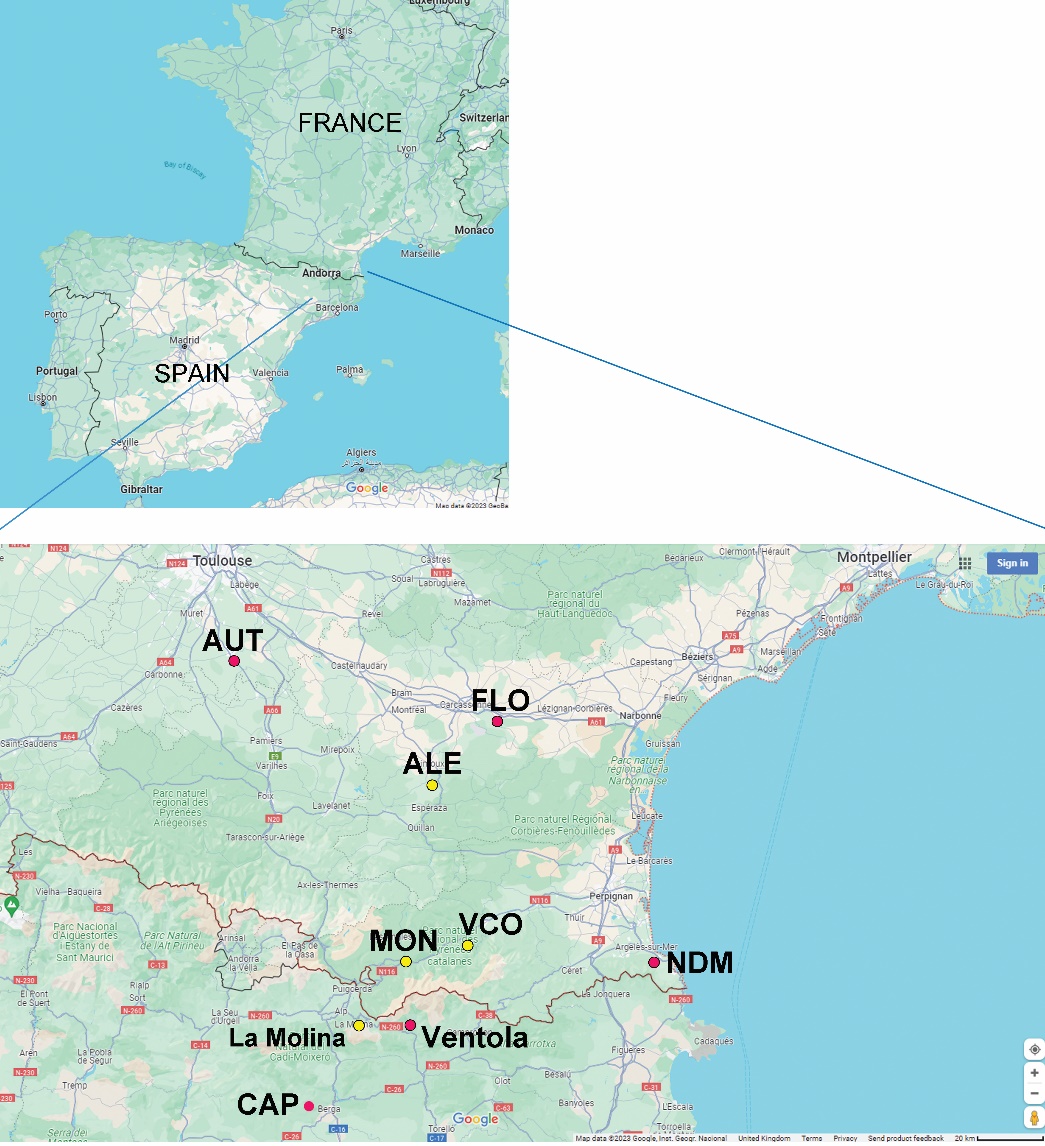

**Supplemental figure S9. Origin of *Antirrhinum* Species Populations Studied.** The wild *Antirrhinum* populations and Accessions used in this study were mapped to a region of south-west Europe. A close-up below, showed their distribution near the border between France and Spain. The Hybrid Zone details between La Molina and Ventola have been described (Tavares *et al*., 2018).

**Supplementary Tables**

| ***LOCUS*** | **Marker** | **Marker** | **Other marker** | **Oligo** | **Oligo sequence** | **Oligo Position (A. majus v4)** |
| --- | --- | --- | --- | --- | --- | --- |
|  | **Type** | **Name** | **Name(s)** | **Name** |  |  |
| ***David's Cline oligos*** | |  |  |  |  |  |
| ***?*** | **KASP** |  | **s1187_290152** |  |  |  |
| ***FLA*** | **KASP** |  | **s316_93292** |  |  |  |
| ***FLA*** | **KASP** |  | **s316_257789** |  |  |  |
| ***SULF*** | **KASP** |  | **s91_39699** |  |  |  |
| ***?*** | **KASP** |  | **s261_720757** |  |  |  |
| ***ROS*** | **KASP** |  | **ros_assembly_543443** | |  |  |
| ***FLAVIA*** | KASP | s316-3 | 5'-FLA prom | #2205 | GATTCCTCAAGCAGAAAACG | Chr2:53,661,809-53,661,828 |
| ***(FLA)*** |  |  | s316_26294 | #2206 | GATTCCTCAAGCAGAAAACA | Chr2:53,661,809-53,661,828 |
|  |  |  | SNP2A | #2207 | GGAGTGCATCCCTGCCGCG | Chr2:53,661,872-53,661,854 |
|  | KASP | s316-R2 | 3'-FLA down | do253 | TTCACGTTCTACGAAGGGGTA | Chr2:53,437,564-53,437,544 |
|  |  |  | s316_250709 | do254 | TTCACGTTCTACGAAGGGGTT | Chr2:53,437,564-53,437,544 |
|  |  |  |  | do255 | CTTTGCCCGTTGCTTGAC | Chr2:53,437,513-53,437,530 |
|  | SANGER |  |  | do259 | TGTTATACGTTTGCGACTCACGAGC | Chr2:53,467,866-53,467,890 |
|  |  |  |  | do460 | GCAGAAGAACAAATTCATCTCCG | Chr2:53,470,017-53,469,995 |
| ***SULFUREA*** | KASP | set 65 | s91_122,561 | do514 | GCAAAATCTGCCCTTTTCCAACTT | Chr4:38,397,156-38,397,179 |
| ***(SULF)*** |  |  |  | do515 | GCAAAATCTGCCCTTTTCCAACTA | Chr4:38,397,156-38,397,179 |
|  |  |  |  | do516 | ACTGATGTGAGCGCCGACTGAGC | Chr4:38,397,207-38,397,185 |
|  | KASP | set 66 | s91_181,717 | do517 | GAATACCACTAAACGAGTGAATGA | Chr4:38,456,336-38,456,359 |
|  |  |  |  | do518 | GAATACCACTAAACGAGTGAATGG | Chr4:38,456,336-38,456,359 |
|  |  |  |  | do519 | CTGAATGTCTTCGAAAGGACAGTG | Chr4:38,456,396-38,456,374 |
| ***AURINA*** | KASP | set 61 |  | do502 | TGGAGTCTTAGCGCTCGACACC | CHr2:1,052,509-1,052,488 |
| ***(AUR)*** |  |  |  | do503 | TGGAGTCTTAGCGCTCGACACA | CHr2:1,052,509-1,052,488 |
|  |  |  |  | do504 | CAATACCACTACTCCTGAAGAGC | Chr2:1,052,428-1,052,450 |
| ***CREMOSA*** | KASP | set 54 B |  | do467 | GTGACTTGGGAGGAAGAATAATC | Chr1:955,163-955,185 |
| ***(CRE)*** |  |  |  | do468 | GTGACTTGGGAGGAAGAATAATA | Chr1:955,163-955,185 |
|  |  |  |  | do469 | TTGGTGATTAAGGGGAAAGTGAC | Chr1:955,225-955,203 |
|  | AFLP | 475-6 |  | do475 | GAGGCTAGGAAGAAAGGTTTGTCG | Chr1:954,562-954,585 |
|  |  |  |  | do476 | CTAACATTGAGCCAAATATTTGCC | Chr1:955,658-955,635 |
| ***ROS1*** | KASP | ROS1 int |  | #1911 | CAACATTGACGTACGGTATTC | Chr6:52,887,667-52,887,647 |
|  |  |  |  | #1912 | CAACATTGACGTACGGTATTT | Chr6:52,887,667-52,887,647 |
|  |  |  |  | #1483 | TGGCATCAAGTTCCACACAGAGCAG | Chr6:52,887,481-52,887,505 |
| ***ELUTA*** | AFLP | 1615-6 |  | #1615 | CATTGTCATGACTCGTTCAACA | Chr6:53,063,047-53,063,068 |
|  |  |  |  | #1616 | TTAAACTGAAAGGCAGGCAATC | Chr6:53,063,517-53,063,496 |
| ***recombination analysis*** | | |  |  |  |  |
| ***ps x str*** | KASP | Fla_set 23 | | do275 | ACTTGGTTGTCCGCGAGAAGCT | Chr2:12,126,989-12,126,968 |
|  |  |  |  | do276 | ACTTGGTTGTCCGCGAGAAGCC | Chr2:12,126,989-12,126,968 |
|  |  |  |  | do277 | CACGGATAAAAAGCAAATCTGGC | Chr2:12,126,933-12,126,955 |
|  | KASP | Fla_set 36 | | do314 | AATCACCGCATTTCTGGAAGCC | Chr2:46,439,115-46,439,094 |
|  |  |  |  | do315 | AATCACCGCATTTCTGGAAGCT | Chr2:46,439,115-46,439,094 |
|  |  |  |  | do316 | TGATGTTTGGACCTTTCTGAGCC | Chr2:46,439,065-46,439,087 |
|  | KASP | Fla_set 33 | | do305 | GAGAAACATAAGCCGAGTGGCC | Chr2:53,067,851-53,067,872 |
|  |  |  |  | do306 | GAGAAACATAAGCCGAGTGGCT | Chr2:53,067,851-53,067,872 |
|  |  |  |  | do307 | TCTCCATGCTGAAACACACGTCG | Chr2:53,067,909-53,067,887 |
|  | KASP | Fla_set 43 | | do364 | GATTTGGTTGAAGAAAACAGTGC | Chr2:53,469,477-53,469,499 |
|  |  |  |  | do365 | GATTTGGTTGAAGAAAACAGTGA | Chr2:53,469,477-53,469,499 |
|  |  |  |  | do366 | AAGCTTCTATATAATTAAGACGG | Chr2:53,469,528-53,469,506 |
|  | KASP | Fla_set 31 | | do299 | GAAGCTGGAGGTTTAGCTCGAT | Chr2:53,594,752-53,594,773 |
|  |  |  |  | do300 | GAAGCTGGAGGTTTAGCTCGAC | Chr2:53,594,752-53,594,773 |
|  |  |  |  | do301 | GAGATTTGCTATCTGGTATTGGAG | Chr2:53,594,824-53,594,801 |
|  | KASP | Fla_set 42 | | do361 | TTACACAACGTTTGTTCATACA | Chr2:53,711,760-53,711,739 |
|  |  |  |  | do362 | TTACACAACGTTTGTTCATACT | Chr2:53,711,760-53,711,739 |
|  |  |  |  | do363 | GTCATGTGATACTTGTTGATCTGC | Chr2:53,711,709-53,711,732 |
|  | KASP | Fla_set 44 | | do367 | ATAGAAGAGGCCCTTACCTAGG | Chr2:54,657,411-54,657,432 |
|  |  |  |  | do368 | ATAGAAGAGGCCCTTACCTAGA | Chr2:54,657,411-54,657,432 |
|  |  |  |  | do369 | AAAGTCCTAAGATACCTCAAAGC | Chr2:54,657,458-54,657,436 |
|  | KASP | Fla_set 30 | | do296 | CACTTTGCCTTCTTTGAATATG | Chr2:59,512,733-59,512,712 |
|  |  |  |  | do297 | CACTTTGCCTTCTTTGAATATA | Chr2:59,512,733-59,512,712 |
|  |  |  |  | do298 | GCTCGAAAGGTTTAGTGATTGTC | Chr2:59,512,677-59,512,699 |
|  | KASP | Fla_set 29 | | do293 | ATCCTTCATAGCTGATGAAATA | Chr2:63,046,469-63,046,448 |
|  |  |  |  | do294 | ATCCTTCATAGCTGATGAAATG | Chr2:63,046,469-63,046,448 |
|  |  |  |  | do295 | AGCTCCGAGCTGAATATGGAGCC | Chr2:63,046,414-63,046,436 |
|  | KASP | Fla_set 38 | | do320 | ATCAAGTATATAACGAGATTGG | Chr2:72,179,984-72,179,963 |
|  |  |  |  | do321 | ATCAAGTATATAACGAGATTGA | Chr2:72,179,984-72,179,963 |
|  |  |  |  | do322 | TGGATCAGGAGAAGTATGTTC | Chr2:72,179,933-72,179,953 |
| ***str x maj*** | KASP | Fla_set 22 | | do272 | GGAAATGTTGTGCAACATTGGC | Chr2:5,356,724-5,356,745 |
|  |  |  |  | do273 | GGAAATGTTGTGCAACATTGGG | Chr2:5,356,724-5,356,745 |
|  |  |  |  | do274 | GGATCTTCATGGAAGGGCAACTC | Chr2:5,356,771-5,356,749 |
|  | KASP | Fla_set 26 | | do284 | GCTCTCAAGTTTCAACCACACG | Chr2:41,257,084-41,257,063 |
|  |  |  |  | do285 | GCTCTCAAGTTTCAACCACACA | Chr2:41,257,084-41,257,063 |
|  |  |  |  | do286 | GTGTGAGGCAAGTTTCAAAAGAG | Chr2:41,257,026-41,257,048 |
|  | KASP | Fla_set 40 | | do326 | AAAATACCGGGGAGAGAGAAGT | Chr2:57,901,885-57,901,906 |
|  |  |  |  | do327 | AAAATACCGGGGAGAGAGAAGC | Chr2:57,901,885-57,901,906 |
|  |  |  |  | do328 | CCTTCTCAATTGAAACCTTCACC | Chr2:57,901,930-57,901,908 |
|  | KASP | Fla_set 28 | | do290 | GCAAATCTTGTTGTACTAACAAC | Chr2:65,601,925-65,601,947 |
|  |  |  |  | do291 | GCAAATCTTGTTGTACTAACAAA | Chr2:65,601,925-65,601,947 |
|  |  |  |  | do292 | GAAGAGATTGAATGGGATTTGCAG | Chr2:65,601,975-65,601,952 |
| ***ps x maj*** | KASP | Fla_set 37 | | do317 | TGAGGTAAGAATGTGCACTGAG | Chr2:29,734,621-29,734,642 |
|  |  |  |  | do318 | TGAGGTAAGAATGTGCACTGAT | Chr2:29,734,621-29,734,642 |
|  |  |  |  | do319 | TATGAATCTTTATCATGACCAAC | Chr2:29,734,673-29,734,651 |
|  | KASP | Fla_set 39 | | do323 | CCATTACTTGAATCAAAGTAAGA | Chr2:64,037,674-64,037,652 |
|  |  |  |  | do324 | CCATTACTTGAATCAAAGTAAGG | Chr2:64,037,674-64,037,652 |
|  |  |  |  | do325 | GTATTCTTCAGCATTGCTTACTC | Chr2:64,037,614-64,037,636 |
|  | KASP | Fla_set 23 | | details as above | |  |
|  | KASP | Fla_set 31 | | details as above | |  |
|  | KASP | Fla_set 36 | | details as above | |  |
|  | KASP | Fla_set 33 | | details as above | |  |
|  | KASP | Fla_set 38 | | details as above | |  |
|  | KASP | Fla_set 42 | | details as above | |  |
|  | KASP | Fla_set 43 | | details as above | |  |

**Supplementary Table S1. Oligos used in this study.**

**Supplemental References (28-31)**

28.	Transposable Element Tam3 at 2 Unlinked Pigment Loci in Antirrhinum-Majus. Molecular & General Genetics 207, 82-89 (1987).

29.	E. S. Coen, R. Carpenter, C. Martin, Transposable Elements Generate Novel Spatial Patterns of Gene-Expression in Antirrhinum Majus. Cell 47, 285-296 (1986).

30.	A. B. Rebocho, P. Southam, J. R. Kennaway, J. A. Bangham, E. Coen, Generation of shape complexity through tissue conflict resolution. Elife 6, (2017).

31.	H. Ringbauer, A. Kolesnikov, D. L. Field, N. H. Barton, Estimating Barriers to Gene Flow from Distorted Isolation-by-Distance Patterns. Genetics 208, 1231-1245 (2018).
